# Evolutionary dynamics of *Aegilops* revealed through comparative genome assembly of all 25 species

**DOI:** 10.64898/2026.07.09.737531

**Authors:** Hamna Shazadee, Tara Edwards, Madeleine Lévesque-Lemay, Chunfang Zheng, Jennifer Ens, Curtis J. Pozniak, Frank M. You, Sylvie Cloutier

## Abstract

*Aegilops* species are the closest wild relatives of wheat and an important reservoir of genetic diversity for its improvement. Despite their potential, many *Aegilops* genomes remain poorly characterized. Here we present high-quality assemblies of 18 diploid, tetraploid, and hexaploid *Aegilops* genomes, which, along with the previously published genomes, complete the production of reference assemblies for all 25 genomes in this genus. Assembly sizes ranged from 5.24 Gb in diploids to 12.65 Gb in hexaploids, with scaffold N50 values up to 749.2 Mb. Gene annotation identified 53,035-156,779 protein-coding genes, of which 21,865-60,490 were classified as high-confidence. Orthogroup-based pangenome analysis across the 25 *Aegilops* genomes identified 80,521 orthogroups, including 15,809 core, 61,735 dispensable, and 2,977 species-specific orthogroups, highlighting substantial gene content variation among genomes. Phylogenetic analysis of 63 *Triticum* and *Aegilops* genomes/subgenomes based on near single-copy orthologs defines the phylogenetic relationships within the *Triticum/Aegilops* complex and confirms diploid progenitors of polyploid lineages. *Ae. mutica* (T) and *Ae. speltoides* (S) belong to the B lineage while the remaining Sitopsis grouped within the D lineage. Structural variation analyses using diploid progenitors as references revealed extensive large-scale rearrangements following polyploidization, emphasizing the dynamics of their evolution. Transposable element (TE) annotation further highlighted subgenome-specific TE expansions and contractions, providing insights into the mechanisms shaping genome structure after polyploidization. Collectively, these genomic resources provide a comprehensive framework for exploring *Aegilops* diversity, understanding polyploid evolution, and accelerating wheat improvement.

## Introduction

Bread wheat (*Triticum aestivum* L.) is a major staple crop, providing approximately 20% of global calories and proteins of the Human population (Gustafsson et al., 2009). As the human population continues to grow, wheat production must increase by more than 50% over current levels by 2050 to meet future food demands (Tadesse et al., 2019). However, increasing productivity while maintaining food safety poses a significant challenge to scientists as agriculture simultaneously faces climate change, evolving pathogens, and environmental stressors. Wheat’s genetic diversity has been reduced by successive bottlenecks during its evolution, especially through polyploidization, domestication, and intensive modern breeding. Overcoming these limitations may be key to addressing major challenges in wheat production (Cavanagh et al., 2013; Charmet, 2011; Dubcovsky and Dvorak, 2007)

Wild relatives, particularly *Aegilops* species, offer a rich source of genetic variation (Dempewolf et al., 2017; Schneider et al., 2008). The genus includes 25 species across eleven genome types (D, S, S^b^, S^l^, S^sh^, S^s^, U, C, N, M and T), with polyploids arising via interspecific hybridization and chromosome doubling (Kilian et al., 2011; Van Slageren, 1994). Following polyploidization, genomes undergo structural changes, gene expansion or loss, and subgenome divergence, producing genomic structures distinct from diploid progenitors despite sharing genome labels (Kilian et al., 2011; Van Slageren, 1994). While phylogenetic relationships have been studied using morphological and molecular data (Dvorak et al., 1998; Dvorak et al., 1993; Petersen et al., 2006; Van Slageren, 1994), genome-scale analyses of structural variation and polyploid evolution remain limited. *Aegilops* species have played a pivotal role in wheat evolution, with *Ae. tauschii* donating the D subgenome (Kihara, 1944; McFadden and Sears, 1946; Salamini et al., 2002), and a species related to *Ae. speltoides* contributing the B subgenome (Li et al., 2022; Marcussen et al., 2014). They also provide valuable alleles for disease resistance and stress tolerance (Colmer et al., 2006; Helguera et al., 2003; Nevo and Chen, 2010; Olivera et al., 2018; Weidner et al., 2012), though their use in breeding is limited by crossing barriers and linkage drag.

Transposable elements (TEs), especially long-terminal repeat (LTR) retrotransposons (*Copia* and *Gypsy*), dominate large grass genomes and contribute to genome size variation, structural remodeling, and gene regulation (Bennetzen and Wang, 2014; Feschotte et al., 2002; Wicker et al., 2018; Wicker et al., 2007). TE dynamics often shift following polyploidization due to genomic shock, relaxed selection, or epigenetic reprogramming (Bennetzen and Wang, 2014; de Tomás and Vicient, 2024; Vicient and Casacuberta, 2017). Despite their importance, the role of TEs in subgenome divergence across *Aegilops* has not been studied because genome assemblies were not available for all species.

The genome assembly of several diploid (7 chromosomes) species have been sequenced, including *Ae. tauschii* (D), *Ae. speltoides* (S), *Ae. longissima* (S^l^), *Ae. bicornis* (S^b^), *Ae. searsii* (S^s^), *Ae. sharonensis* (S^sh^), *Ae. umbellulata* (U), *Ae. comosa* (M), and *Ae. mutica* (T) (Abrouk et al., 2023; Avni et al., 2022; Grewal et al., 2025; Li et al., 2024; Li et al., 2022; Luo et al., 2017; Wang et al., 2021). Our lab has recently completed the assemblies of the remaining diploid species, including *Ae. uniaristata* (N) and *Ae. markgrafii* (C) (Khadka et al. 2026, unpublished). However, the only genome assembly of polyploid (14 and 21 chromosomes) species published to date is that of *Ae. ventricosa* (DN) (Liu et al., 2025), leaving a knowledge gap in polyploid *Aegilops* structural genomics.

To address this gap, we generated high-quality assemblies for 18 *Aegilops* species (4 diploid, 10 tetraploid and 4 hexaploid) using Oxford Nanopore, PacBio, Illumina, Hi-C, and Omni-C technologies. Consistent machine learning–based gene annotation, phylogenomic analyses, structural variation, and LTR characterization enabled a comprehensive view of subgenome evolution, TE dynamics, and divergence from diploid progenitors. Collectively, this genomic resource provides a foundation for understanding *Aegilops* genome evolution and exploiting wild diversity for wheat improvement.

## Results

### Genome Assembly

*Aegilops* species exhibit substantial morphological diversity at the whole-plant level, reflecting their wide evolutionary range across diploid, tetraploid, and hexaploid lineages (Figure 1A). To investigate the genomic basis underlying this diversity, we generated high-quality genome assemblies for 18 *Aegilops* species, representing four of the 11 diploid, all ten tetraploid, and all four hexaploid species, using a single representative accession for each species (Supplemental Table 1). Assemblies were produced using Oxford Nanopore Technologies (ONT, Oxford, UK) long reads, polished with PacBio (Pacific Biosciences, Menlo Park, CA, USA) HiFi long reads and Illumina (San Diego, CA, USA) short reads, and scaffolding with Hi-C and Omni-C (Dovetail Genomics, Scotts Valley, CA, USA) short-read data. Sequencing coverage for each platform and genome assembly is summarized in Supplemental Table 2. Diploid genome sizes ranged from 5.24 Gb (*Ae. searsii*) to 6.53 Gb (*Ae. longissima*), with N50 values of 560.0–749.2 Mb and N90 values of 23.2–397.7 Mb (Supplemental Table 3). Scaffold numbers ranged from 226 to 1,720, with 8–10 scaffolds exceeding 100 Mb (Figures 1B, 1C and Supplemental Table 3). Tetraploid genomes ranged from 7.40 Gb (*Ae. cylindrica*) to 9.79 Gb (*Ae. kotschyi*), with N50 values of 367.2–644.9 Mb and N90 values of 62.5–339.7 Mb (Figure 1B and Supplemental Table 3). Scaffold counts ranged from 349 to 2,463, with 14–24 scaffolds >100 Mb (Figures 1B-1C and Supplemental Table 3). Hexaploid assemblies ranged from 11.57 Gb (*Ae. vavilovii*) to 12.65 Gb (*Ae. neglecta* subsp. *recta*), with N50 values of 623.8–705.7 Mb and N90 values of 316.1–390.8 Mb (Figure 1B and Supplemental Table 3). Scaffold numbers ranged from 266 to 1,070, with 21–23 scaffolds >100 Mb (Figures 1B-1C and Supplemental Table 3). Scaffolds >100 Mb accounted for 75.62–98.27% of diploid assemblies, 86.98–99.79% of tetraploid assemblies, and 98.95–99.89% of hexaploid assemblies, whereas scaffolds <100 Mb contributed only 0.66–8.90%, 0.93–3.92%, and 1.78–2.82% of the total assembly size, respectively, illustrating the high contiguity of all genome assemblies.

**Figure 1.**
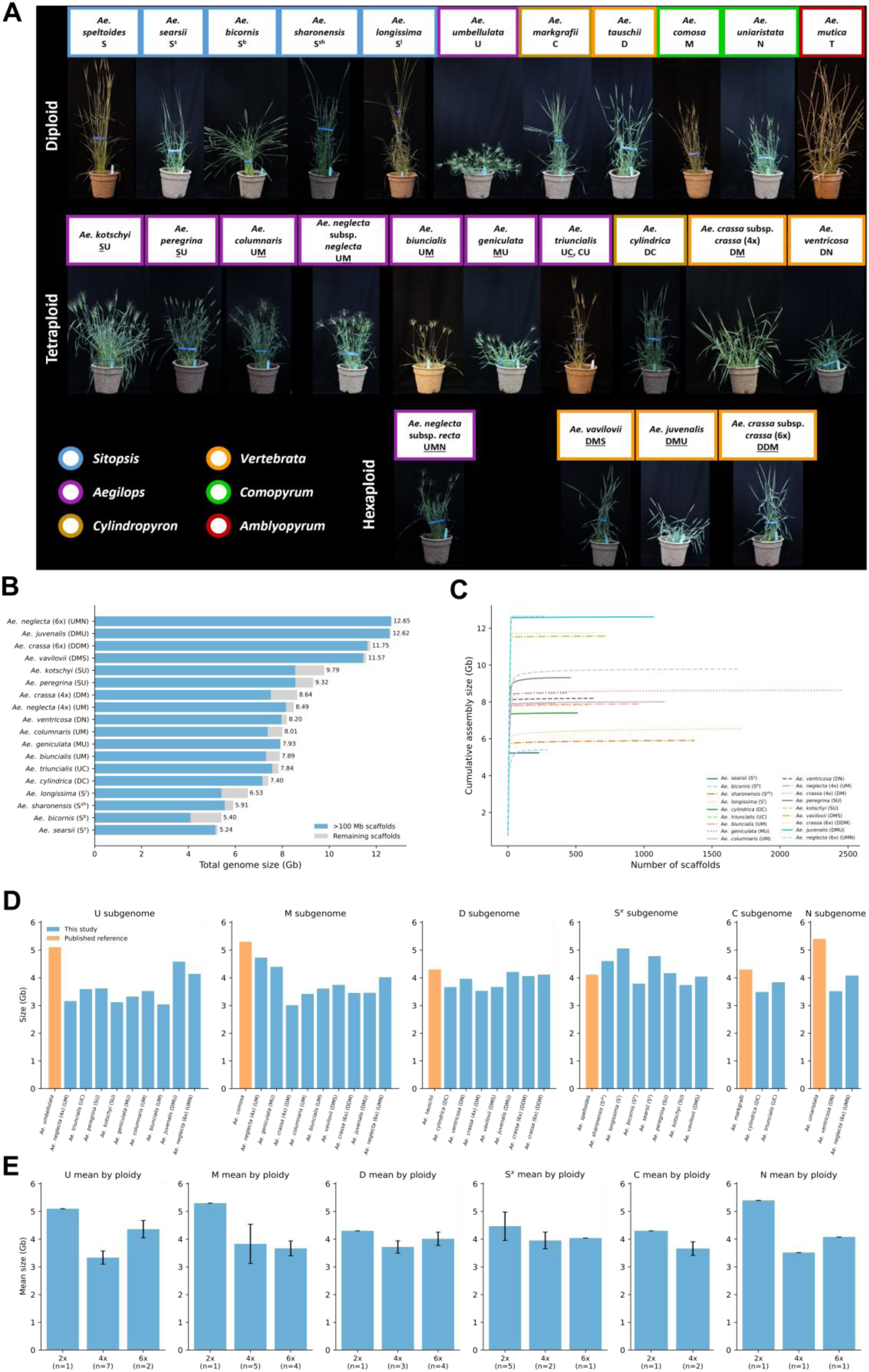
Morphological diversity and genome assembly statistics across *Aegilops* species. **(A)** Whole-plant morphology showing variation in plant architecture across diploid, tetraploid, and hexaploid *Aegilops* species, highlighting extensive phenotypic diversity across the genus. **(B)** Total genome size estimates for each species, with contributions from scaffolds >100 Mb shown alongside remaining scaffolds. **(C)** Assembly contiguity illustrated by the relationship between cumulative assembly size and number of scaffolds across genomes, reflecting variation in scaffold distribution and assembly structure among species. **(D)** Comparative genome size variation across subgenomes (U, M, D, Sˣ, C and N) in polyploid *Aegilops* species relative to their corresponding diploid progenitors, highlighting subgenome-specific expansion and contraction patterns. **(E)** Mean genome size variation across ploidy levels (diploid, tetraploid, and hexaploid) for each subgenome group, showing trends in genome size evolution following polyploidization.

Benchmarking Universal Single-Copy Orthologs (BUSCO) analysis using the embryophyta_odb10 dataset (Manni et al., 2021) showed high completeness (98.4–99.8%), supporting the overall quality of the assemblies (Supplemental Figure 1 and Supplemental Table 3).

Comparative analyses of genome size between polyploid subgenomes and their corresponding diploid reference genomes revealed a consistent reduction in subgenome size following polyploidization, indicating genome contraction relative to their putative progenitors (Figure 1D). A similar trend was observed when comparing mean genome sizes across ploidy levels, where diploid genomes were consistently larger, whereas tetraploid and hexaploid subgenomes exhibited reduced average genome sizes across all genome groups (U, M, D, S^x^, C and N) (Figure 1E). Together, these results demonstrate that the generated assemblies achieved high contiguity and completeness across all ploidy levels while also revealing consistent patterns of genome downsizing associated with polyploid evolution in *Aegilops*.

### Gene annotation and orthogroup-based pangenome analysis

The total numbers of protein-coding genes annotated increased with ploidy, ranging from 53,035–59,817 in diploids, 88,422–109,856 in tetraploids and 143,429–156,779 in hexaploids. *Ae. crassa* subsp. *crassa* (4x) was an outlier with 149,108 genes. HC gene counts ranged from 21,865–27,783 (diploids), 33,590–60,490 (tetraploids), and 44,149–55,945 (hexaploids). BUSCO completeness scores (84.5%–98.9%) indicate that most core genes were captured (Figure 2A and Supplemental Table 4).

**Figure 2.**
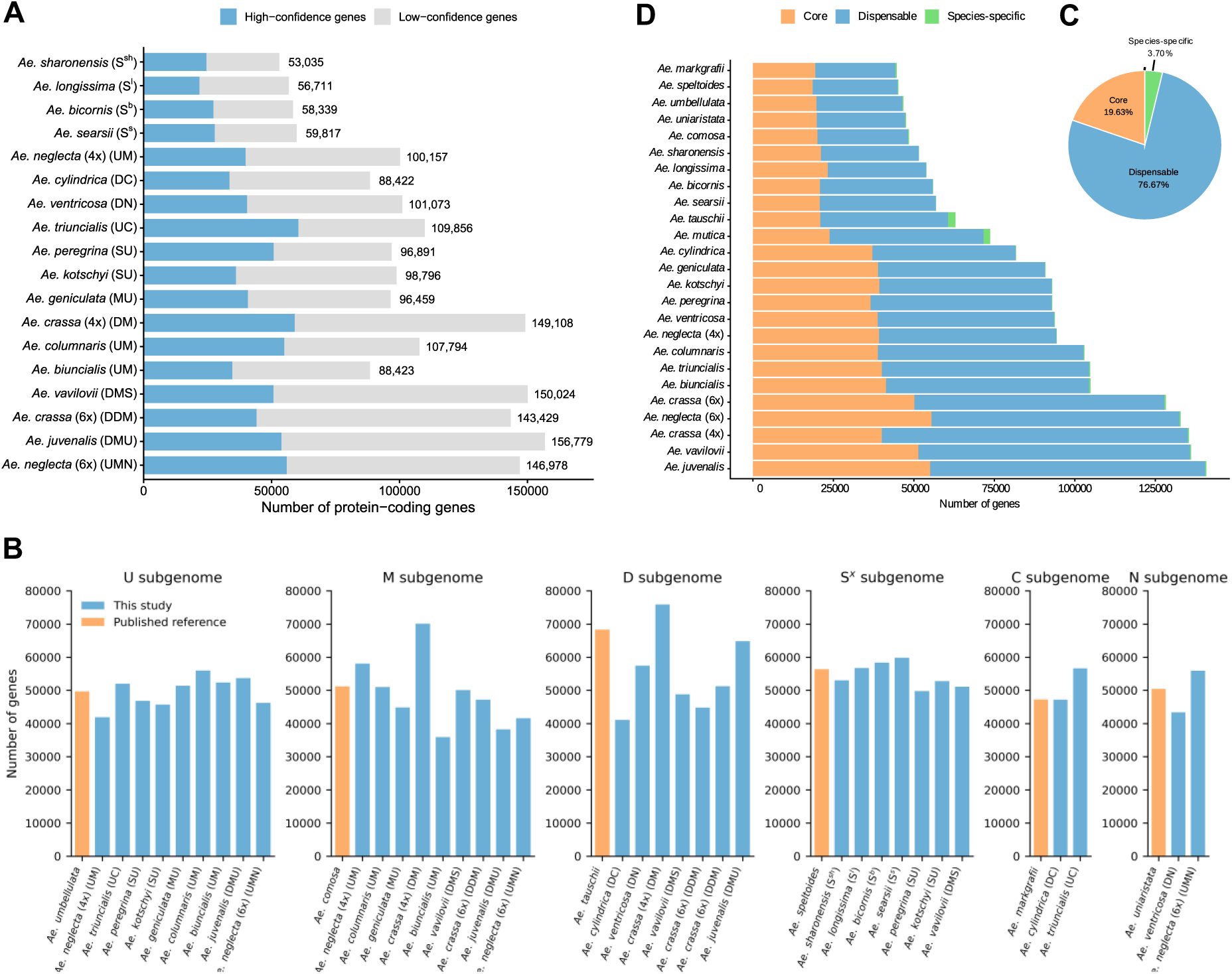
Gene annotation statistics and pan-genome analysis of *Aegilops* species based on protein-coding genes. **(A)** Bar plot showing the total number of protein-coding genes in each species, including high-confidence and low-confidence gene models, highlighting variation in gene content across the genus. **(B)** Comparison of total protein-coding gene numbers across subgenome groups (U, M, D, Sˣ, C and N) relative to their corresponding diploid progenitor species, illustrating changes in gene content associated with polyploidization. **(C)** Pie chart showing the proportions of core, dispensable, and species-specific orthogroups across the *Aegilops* pan-genome, summarizing gene conservation and diversification across species. **(D)** Distribution of core, dispensable, and species-specific genes across individual genomes, highlighting variation in gene sharing and lineage-specific gene content. The following species names were shortened: *Ae. neglecta* subsp. *neglecta* is shown as *Ae. neglecta* (4x); *Ae. neglecta* subsp. *recta* as *Ae. neglecta* (6x); *Ae. crassa* subsp. *crassa* (4x) as *Ae. crassa* (4x); *Ae. Crassa* subsp. *crassa* (6x) as *Ae. crassa* (6x).

To further resolve gene content evolution at the subgenome level, we compared total protein-coding gene numbers in each polyploid subgenome against their corresponding diploid progenitors (Figure 2B). This revealed highly heterogeneous patterns of gene retention and loss across subgenomes following polyploidization. The U subgenome showed both expansion and contraction relative to its diploid reference, indicating lineage-dependent gene content evolution rather than a uniform trend. In contrast, the M subgenome generally exhibited gene loss across most polyploid species, with notable exceptions of gene expansion in *Ae. crassa* subsp. *crassa* (4x) and *Ae. neglecta* subsp. *neglecta*, suggesting pervasive fractionation following polyploidization. The D subgenome displayed the highest variability among all subgenomes. Strong gene expansion was observed exclusively in *Ae. crassa* subsp. *crassa* (4x), whereas all other comparisons showed reduced gene numbers relative to diploid *Ae. tauschii*. This pattern highlights highly dynamic and lineage-specific evolutionary trajectories within the D subgenome. In comparison, the S subgenome remained largely unchanged, with gene contents closely matching its diploid counterpart. Similarly, the C subgenome showed high conservation, with *Ae. cylindrica* maintaining nearly identical gene numbers to diploid *Ae. markgrafii*, while *Ae. triuncialis* exhibited a moderate expansion. The N subgenome showed intermediate behavior, with gene loss in *Ae. ventricosa* and gene expansion in *Ae. neglecta subsp. recta*, indicating moderate but lineage-specific variation without a consistent directional trend (Figure 2B). Overall, these results indicate that gene content evolution in *Aegilops* polyploids is strongly subgenome-dependent, with distinct evolutionary trajectories ranging from high conservation (S and C subgenomes), moderate plasticity (N and U subgenomes), to pronounced structural and gene content remodeling (D and M subgenomes). This subgenome-specific behavior suggests that post-polyploidization genome evolution is not uniform but instead shaped by differential fractionation, gene retention, and lineage-specific genomic restructuring.

Orthogroup-based pangenome analysis further resolved gene content variation across the 25 *Aegilops* genomes. A total of 80,521 orthogroups were identified, including 15,809 (19.63%) core orthogroups shared across all genomes, 61,735 (76.67%) dispensable orthogroups present in 2–24 genomes, and 2,977 (3.70%) species-specific orthogroups, unique to individual genomes where the latter ranged from 16 in *Ae. sharonensis* to 548 in *Ae. mutica* (Figure 2C and Supplemental Table 5). These orthogroups corresponded to 18,587–55,519 core genes, 24,913–95,073 dispensable genes, and 36–2,283 species-specific genes per genome (Figure 2D and Supplemental Table 5) with both core and dispensable gene fractions generally increasing with ploidy level.

### Phylogenetic relationships and divergence time

Phylogenetic relationships were inferred from near single-copy orthologs (773) across 36 *Aegilops* genomes and subgenomes generated in this study, together with 27 previously published *Aegilops* and *Triticum* genomes/subgenomes. *Brachypodium distachyon*, *Hordeum vulgare* and *Thinopyrum elongatum* were included as outgroups to root the tree. The resulting phylogeny resolved well-supported A, B, D, S^x^ (Sitopsis) N, M, C, and U clades (Figure 3). Within the A clade, the A genomes of *T. urartu* and *T. monococcum* clustered with the A subgenomes of polyploid *Triticum* species. The B genome clade included the B subgenomes of *T. aestivum* and *T. turgidum*, the S genome of the *Ae. speltoides*, the T genome of *Ae. mutica* and the G genome of *T. timopheevii*. Most polyploid *Aegilops* subgenomes clustered with their expected diploid progenitors. For example, the D subgenomes grouped with the D genome of *Ae. tauschii* and D subgenome of *Triticum aestivum*, reflecting their shared ancestry. The N subgenomes of *Ae. ventricosa* (DN) and *Ae. neglecta* subsp. *recta* (UMN) grouped with *Ae. uniaristata* (N), while the C subgenomes of *Ae. triuncialis* (UC) and *Ae. cylindrica* (DC) clustered with *Ae. markgrafii* (C). Similarly, all U subgenomes formed a well-supported group with *Ae. umbellulata* (U). The “S” subgenomes of *Ae. kotschyi* and *Ae. peregrina* clustered within the Sitopsis group, close to *Ae. longissima* (S^l^) and *Ae. sharonensis* (S^sh^), hinting that one or the other are the most likely progenitors of these species rather than *Ae. speltoides* (Figure 3). These concordant groupings confirm the involvement of the corresponding diploid species as progenitors of the respective subgenomes in polyploid *Aegilops* species.

**Figure 3.**
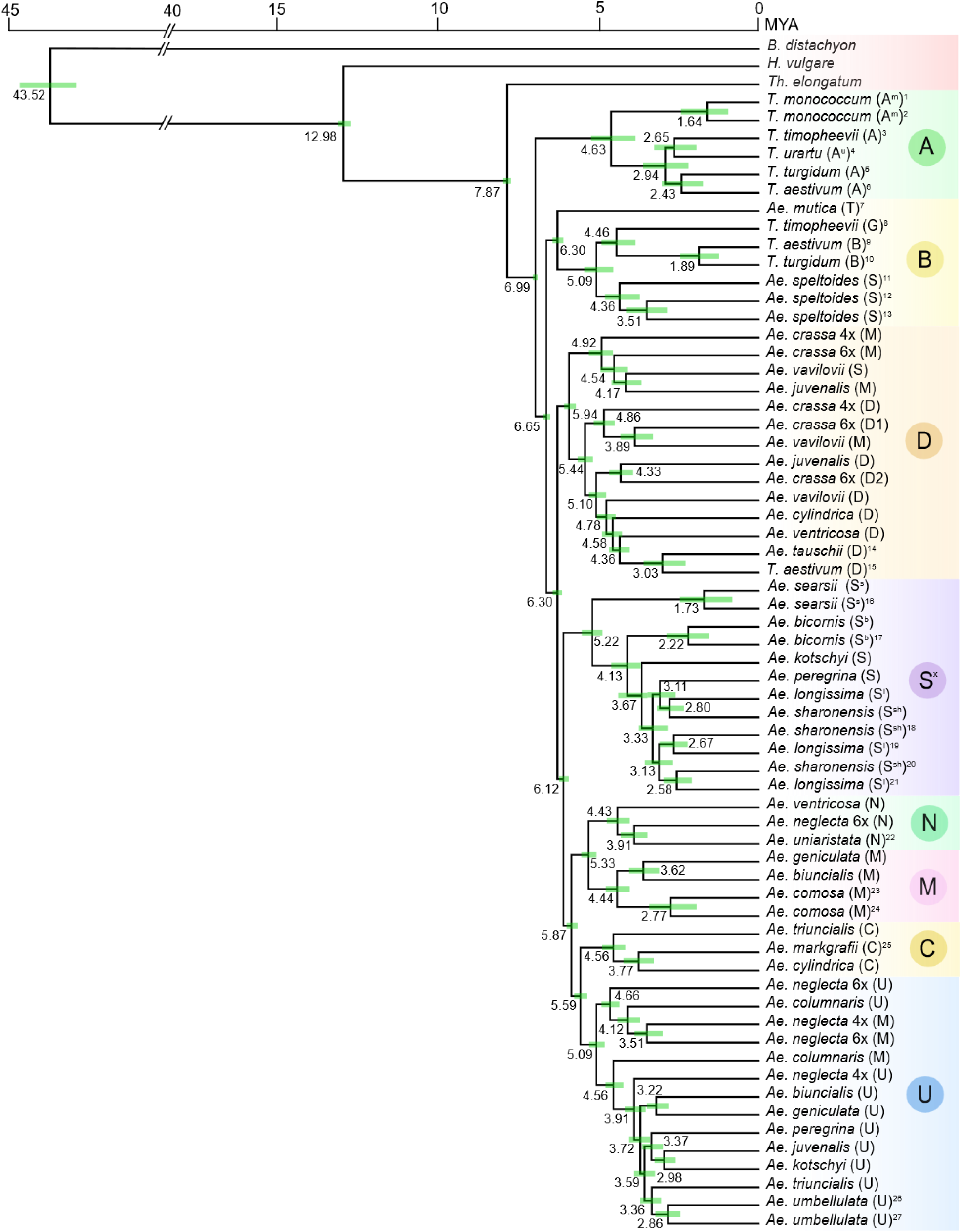
Phylogenetic relationships and divergence-time estimates in the *Triticum*/*Aegilops* species complex. Phylogeny and divergence times inferred from single-copy orthologs. Horizontal bars represent the 95% highest posterior density (HPD) intervals for each node. Superscript numbers correspond to previously published genome references (e.g., 1, 2 — Ahmed et al. (2023); 3, 8 — Grewal et al. (2024); 4 — Ling et al. (2018); 5, 10 — Zhu et al. (2019); 6, 9, 15 — IWGSC (2021); 7 — Grewal et al. (2025); 11 — Yang et al. (2023); 12, 16, 17, 20, 21 — Li et al. (2022); 13, 18, 19 — Avni et al. (2022); 14 — Wang et al. (2021); 22, 24, 25, 27 — Khadka et al. (2026, unpublished); 23 — Li et al. (2024); 26 — Abrouk et al. (2023). Genomes without superscript numbers represent assemblies generated in this study. For simplicity, the following species names were shortened: *Ae. crassa* subsp. *crassa* (4x) is shown as *Ae. crassa* 4x; *Ae. crassa* subsp. *crassa* (6x) as *Ae. crassa* 6x; *Ae. neglecta* subsp. *neglecta* as *Ae. neglecta* 4x; and *Ae. neglecta* subsp. *recta* as *Ae. neglecta* 6x. A, B, D, S^x^, N, M, C, and U indicate the corresponding clades.

Conversely, the M subgenomes were more phylogenetically dispersed. Only the M subgenomes of *Ae. geniculata* (MU) and *Ae. biuncialis* (UM) clustered with *Ae. comosa* (M). The M subgenomes of *Ae. neglecta* subsp*. neglecta* (UM)*, Ae. neglecta* subsp. *recta* (UMN) and *Ae. columnaris* (UM) clustered within the U-genome clade, while that of *Ae. crassa* subsp. *crassa* (4x, DM), *Ae. crassa* subsp. *crassa* (6x, DDM), *Ae. juvenalis* (DMU), and *Ae. vavilovii* (DMS) nested within the D clade (Figure 3). These interspersed patterns suggest that M subgenomes have undergone extensive interchromosomal rearrangements and genome restructuring following polyploidization, resulting in reduced phylogenetic affinity to their putative diploid progenitors and closer association with co-resident subgenomes within the polyploid nucleus. Similarly, the S subgenome of *Ae. vavilovii* clustered with the D clade rather than with *Ae. speltoides* or any other Sitopsis species. Its close relationship to *Ae. crassa* subsp. *crassa* (6x, DDM) supports the possibility that this *Ae. vavilovii* accession may be misclassified.

Using the same phylogenomic dataset, divergence times among *Triticum–Aegilops* lineages revealed a clear temporal structure. The earliest separations between the A and B genomes were estimated at ∼6.99 MYA, followed by the divergence of the B and D lineages at ∼6.65 MYA. Within the D clade, divergence ranged from 3.03 to 5.44 MYA when considering the D genomes of *Ae. tauschii*, *Triticum aestivum*, and polyploid *Aegilops* species. The most recent split occurred between *Ae. tauschii* and *Triticum aestivum*, whereas the deepest divergence reflects separation among more distantly related D subgenomes. The Sitopsis lineage diverged from the D lineage at ∼6.30 MYA, with internal divergence spanning 1.73–5.22 MYA (Figure 3).

The N, M, C, and U genomes share a common ancestor at ∼5.87 MYA. N genomes exhibit relatively narrow divergence (3.91–4.43 MYA), with the N subgenome of *Ae. neglecta* subsp. *recta* diverging at 3.91 MYA and that of *Ae. ventricosa* slightly earlier at 4.43 MYA. Diploid *Ae. comosa* clustered with the M subgenomes of *Ae. biuncialis* and *Ae. geniculata*, which diverged from each other at 3.62 MYA and from *Ae. comosa* at 4.44 MYA. The M subgenomes that grouped within the D or U clades showed divergence times of 3.89–4.92 MYA (D clade) and 3.51–4.56 MYA (U clade), highlighting their deeper and more variable evolutionary placement. In the C clade, the C subgenomes of *Ae. cylindrica* and *Ae. triuncialis* diverged from the diploid *Ae. markgrafii* at approximately 3.77 and 4.56 MYA, respectively. In contrast, U genomes exhibited a broader divergence range (2.86–5.09 MYA), although most U subgenomes diverged more recently (<4 MYA) (Figure 3). Overall, phylogenetic relationships largely reflect expected diploid–polyploid associations, with exceptions—particularly in the M lineage—indicating a complex, reticulate evolutionary history. Divergence estimates further support earlier origins for M-genome–containing species relative to other lineages.

Phylogeny and divergence times inferred from 773 near single-copy orthologs. Horizontal bars represent the 95% highest posterior density (HPD) intervals for each node. Superscript numbers correspond to previously published genome references (e.g., 1, 2 — Ahmed et al. (2023); 3, 8 — Grewal et al. (2024); 4 — Ling et al. (2018); 5, 10 — Zhu et al. (2019); 6, 9, 15 — IWGSC (2021); 7 — Grewal et al. (2025); 11 — Yang et al. (2023); 12, 16, 17, 20, 21 — Li et al. (2022); 13, 18, 19 — Avni et al. (2022); 14 — Wang et al. (2021); 22, 24, 25, 27 — Khadka et al. (2026, unpublished); 23 — Li et al. (2024); 26 — Abrouk et al. (2023). Genomes without superscript numbers represent assemblies generated in this study. For simplicity, the following species names were shortened: *Ae. crassa* subsp. *crassa* (4x) is shown as *Ae. crassa* 4x; *Ae. crassa* subsp. *crassa* (6x) as *Ae. crassa* 6x; *Ae. neglecta* subsp. *neglecta* as *Ae. neglecta* 4x; and *Ae. neglecta* subsp. *recta* as *Ae. neglecta* 6x. A, B, D, S^x^, N, M, C, and U indicate the corresponding clades.

### Genome structural variation (SV) landscape

To investigate structural variation, tetraploid and hexaploid *Aegilops* subgenomes were compared with their respective diploid progenitors from which synteny, inversions, translocations, duplications, insertions, and deletions were tallied. Many subgenome–progenitor pairs retained extensive synteny, reflecting overall genome conservation. Among the 34 comparisons, fourteen were highly syntenic with over 2,000 shared syntenic blocks, ten showed moderate synteny with 1,000–2,000 blocks, and the remaining comparisons had fewer than 1,000 blocks, indicating varying levels of structural divergence (Figure 4A and Supplemental Table 6).

**Figure 4.**
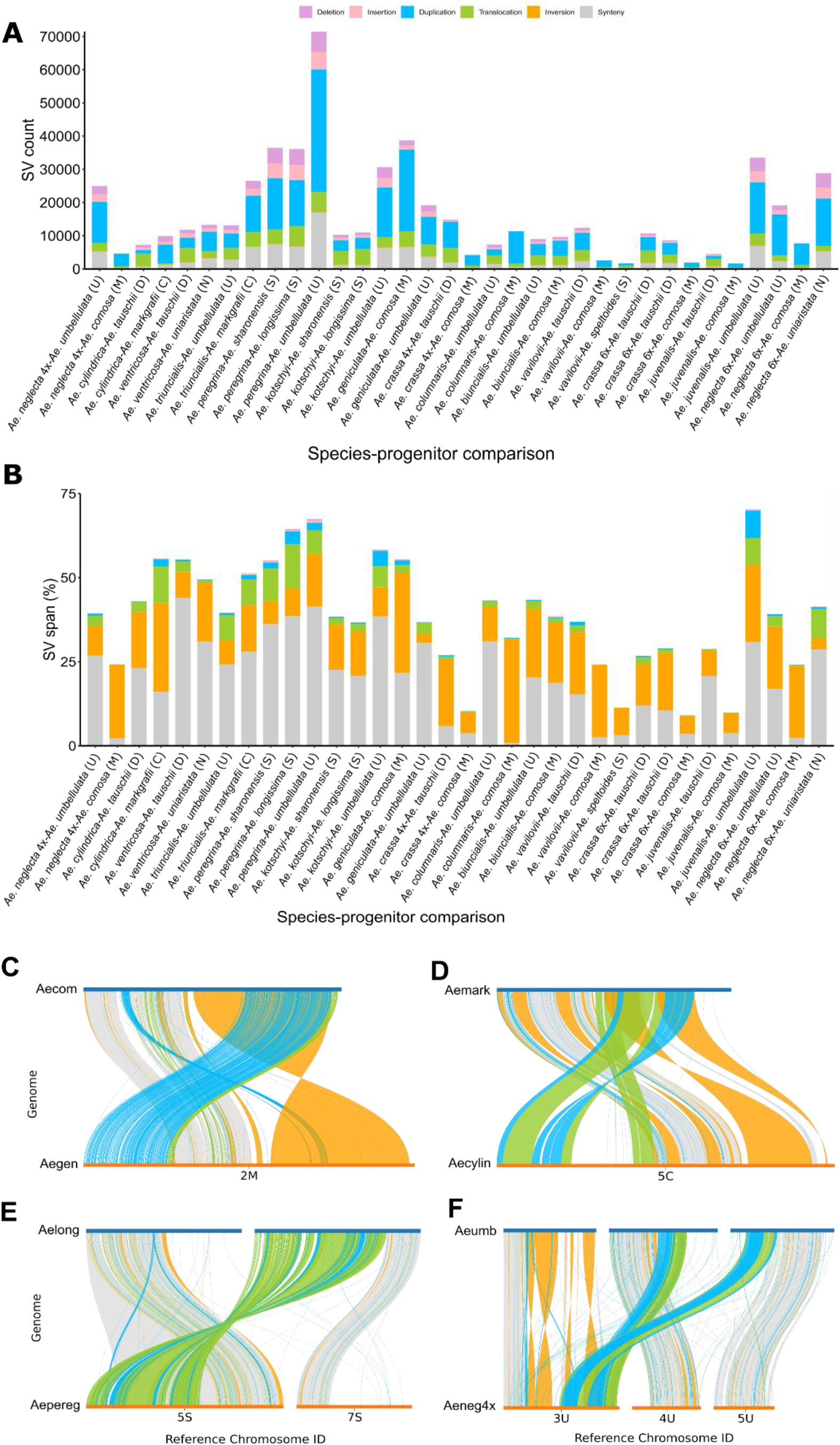
Synteny patterns and structural variation in *Aegilops* subgenomes relative to their diploid progenitors. **(A)** Stacked bar plots showing the distribution of syntenic regions and structural variant (SV) types—including inversions, translocations, duplications, insertions, and deletions—across subgenome–progenitor comparisons. Each bar represents the relative abundance of synteny and SV classes for a given comparison. The following species names were shortened: *Ae. neglecta* subsp. *neglecta* is shown as *Ae. neglecta* 4x; *Ae. crassa* subsp. *crassa* (4x) as *Ae. crassa* 4x; *Ae. crassa* subsp. *crassa* (6x) as *Ae. crassa* 6x; and *Ae. neglecta* subsp. *recta* as *Ae. neglecta* 6x. **(B)** Stacked bar plots showing the percentage of each genome spanned by syntenic regions and SV classes across subgenome–progenitor comparisons. Each bar represents the proportion of genomic sequence covered by each category for a given comparison. Species abbreviations are the same as in (A). **(C–F)** Representative large-scale structural variants identified in polyploid *Aegilops* subgenomes: **(C)** the largest inversion on chromosome 2M in the *Ae. geniculata*–*Ae. comosa* comparison; **(D)** the largest intrachromosomal translocation on chromosome 5C in *Ae. cylindrica*–*Ae. markgrafii*; **(E)** the largest interchromosomal translocation involving chromosome 5S of *Ae. peregrina* and chromosome 7S of *Ae. longissima*; **(F)** the largest interchromosomal duplications between 3U–5U and 3U–4U in *Ae. neglecta* subsp. *neglecta*–*Ae. umbellulata*. To improve visualization, only SVs larger than 10 kb were included in plots. Species abbreviations used in the plots are: Aecom (*Ae. comosa*), Aegen (*Ae. geniculata*), Aemark (*Ae. markgrafii*), Aecylin (*Ae. cylindrica*), Aelong (*Ae. longissima*), Aepereg (*Ae. peregrina*), Aeumb (*Ae. umbellulata*) and Aeneg4x (*Ae. neglecta* subsp. *neglecta*).

Inversions were less frequent than other SV types (Supplemental Figure 2) but often spanned large regions ranging from 507 bp to 351 Mb, and accounted for the largest proportion of genome span among all SV classes (2.6–30.9%) (Figure 4B, Supplemental Tables 6 and 7), with the largest detected on chromosome 2M in the *Ae. geniculata*–*Ae. comosa* comparison (Figure 4C). Translocations were widespread, ranging from 439 bp to 67 Mb, and accounted for 0.0–13.2% of the genome span, with most comparisons showing over 2,000 events (Supplemental Figure 2, Supplemental Tables 6 and 7). The largest intrachromosomal translocation occurred on chromosome 5C in *Ae. cylindrica*–*Ae. markgrafii* (Figure 4D) while the largest interchromosomal event involved chromosomes 5S of *Ae. peregrina* and 7S^l^ of *Ae. longissima* (Figure 4E). Duplications were the most abundant SV type, with ten comparisons exceeding 10,000 events and up to 36,853 in the U genome comparison between *Ae. peregrina* and *Ae. umbellulate* (Figure 4A). Despite this high abundance, duplications accounted for only a small proportion of genome span (0.0–8.2%) (Supplemental Table 6). Duplication sizes ranged from 397 bp to 100 Mb (Supplemental Table 7), with the largest events observed between chromosomes 3U–5U (100 Mb) and 3U–4U (80 Mb) in *Ae. neglecta* subsp. *neglecta* and *Ae. umbellulata* (Figure 4F). Insertions and deletions were less frequent, generally below 2,000 events per comparison, and both spanned only a very small portion of the genome, with sizes ranging from 51 bp to 11 Mb and 51 bp to 2 Mb, respectively (Supplemental Tables 6 and 7).

Lineage-specific patterns were also evident. Several M subgenomes showed low synteny with *Ae. comosa*, resulting in few detectable SVs, consistent with their phylogenetic placement alongside co-resident U or D subgenomes. Subsequent SV analyses of these subgenomes using the reference genome corresponding to their coexisting counterparts revealed higher synteny and more SVs (Supplemental Figures 3A-3G and Supplemental Table 8), indicating extensive post-polyploidization restructuring. Similarly, the putative S subgenome of *Ae. vavilovii* showed limited synteny with *Ae. speltoides* (Supplemental Figure 2Y) and other Sitopsis genomes, but higher synteny with *Ae. tauschii* (Supplemental Figures 3H-3L and Supplemental Table 8). In agreement with its phylogenetic placement, this SV pattern supports the interpretation that this subgenome corresponds to a D genome related to *Ae. crassa* subsp. *crassa* (6x, DDM), indicating potential misclassification of this accession. In contrast, the S subgenomes of *Ae. peregrina* and *Ae. kotschyi* exhibited greater synteny with *Ae. longissima* and *Ae. sharonensis* (Supplemental Figures 2I-2J and 2L-2M) than with *Ae. speltoides* (Supplemental Figures 3M-3N and Supplemental Table 8), consistent with their phylogenetic relationships. Together, SV patterns closely mirror phylogenetic relationships and highlight extensive interchromosomal restructuring following polyploidization.

Structural variation analyses were performed between subgenomes within each polyploid species to assess the extent of genomic differentiation among constituent genomes. Most subgenome pairs exhibited relatively low levels of synteny, indicating substantial divergence and the retention of distinct structural architectures despite coexisting within the same nucleus (Supplemental Figure 4; Supplemental Table 9). In contrast, the U and C subgenomes of *Ae. triuncialis*, the D and M subgenomes of *Ae. crassa* subsp. *crassa* (4x), and the U and M subgenomes of *Ae. columnaris* exhibited comparatively high levels of synteny, with 1,289, 500, and 684 syntenic blocks identified, respectively. These same comparisons also contained large numbers of structural variants. The U and C subgenomes of *Ae. triuncialis* contained 7,535 translocations and 67,758 duplications, the D and M subgenomes of *Ae. crassa* subsp. *crassa* (4x) contained 3,898 translocations and 12,459 duplications, and the U and M subgenomes of *Ae. columnaris* contained 3,220 translocations and 23,725 duplications (Supplemental Figure 4; Supplemental Table 9). Together, these results suggest that although these subgenome pairs retain substantial collinearity and shared genomic content, they have also experienced extensive structural reorganization. Overall, the degree of subgenome differentiation varies considerably among polyploid *Aegilops* species, with most genomic combinations exhibiting substantial divergence, whereas *Ae. triuncialis*, *Ae. crassa* subsp. *crassa* (4x), and *Ae. columnaris* retain relatively high levels of collinearity.

To assert that the extent of the variation observed in polyploid species compared to their progenitor was the result of post-polyploidization restructuring, we quantified SV in four diploid species where assemblies for multiple accessions were available, namely *Ae. sharonensis, Ae. longissima*, *Ae. bicornis* and *Ae. searsii*. Overall, these pan-genome comparisons were more syntenic but more SV could be detected (Supplemental Figure 5 and Supplemental Table 10). This pattern is consistent with differences in sequence similarity between comparison pairs, where higher similarity among diploid genomes enables more extensive alignment and increased SV detection, whereas greater divergence in polyploid–diploid comparisons reduces regions that can be aligned and limits SV detection. Overall, structural variation is pervasive both within diploid *Aegilops* species and in polyploid–diploid comparisons, with the extent of detected variation strongly influenced by the level of genomic similarity between genomes.

### Genome-wide transposable element (TE) composition

Genome-wide transposable element (TE) content was analyzed across all 18 *Aegilops* assemblies. TEs account for 85.44–88.10% of the genomes. This repetitive fraction is predominantly composed of Class I LTR retrotransposons (*Gypsy* and *Copia*) and Class II DNA transposons of the CACTA superfamily, whereas other TE superfamilies contribute marginally. *Gypsy* elements are the most abundant (38.0–46.84%), followed by *Copia* (13.97–19.04%) and CACTA elements (8.54–16.01%) (Figure 5A and Supplemental Table 11).

**Figure 5.**
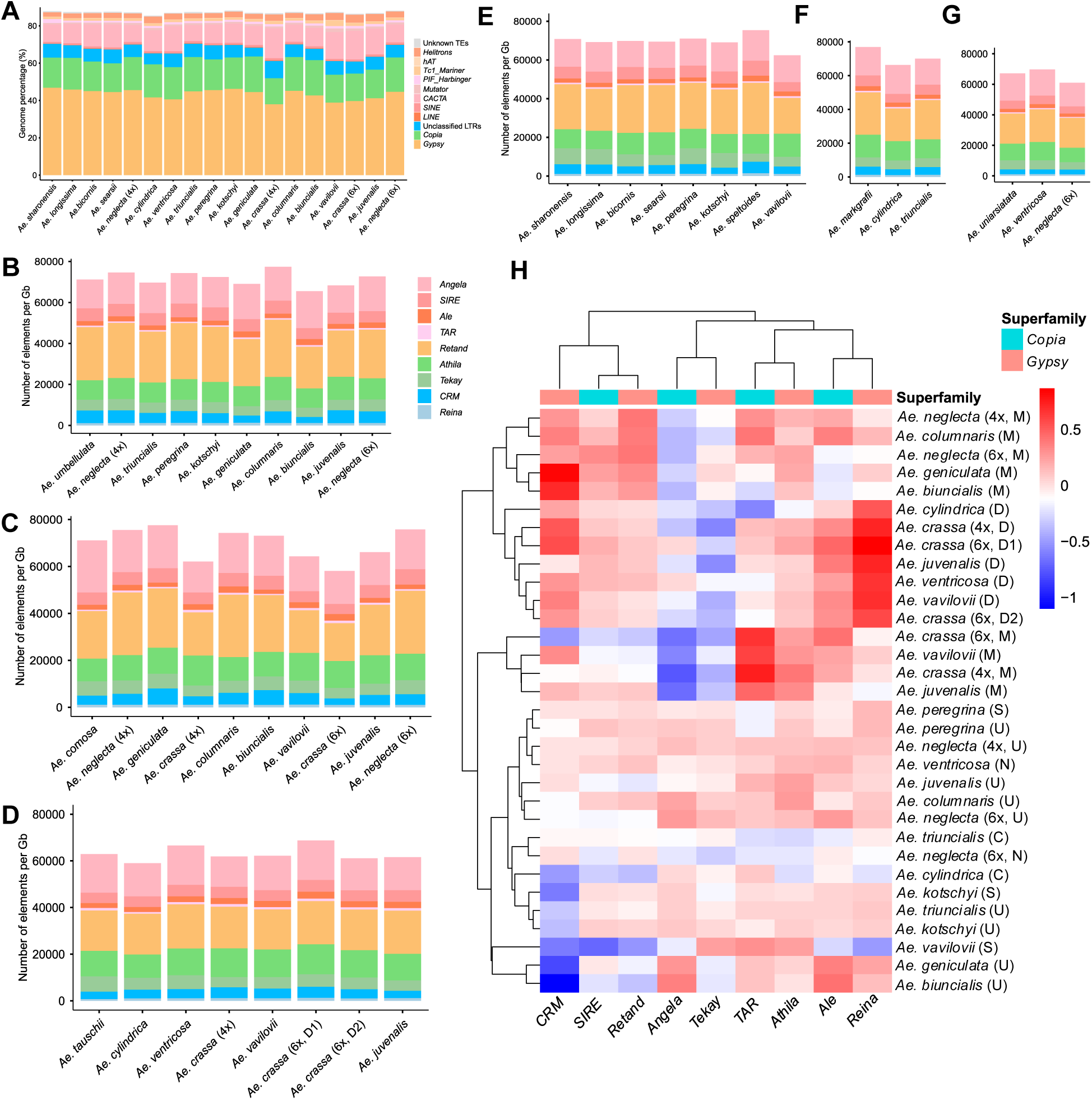
Transposable element composition and LTR clade dynamics across *Aegilops* genomes and subgenomes. **(A)** Stacked bar plot showing the proportional contribution of each transposable element class to overall *Aegilops* genome composition. **(B–G)** Normalized LTR lineage abundance (per Gb) across U, M, D, S, C, and N subgenomes, respectively. **(H)** Heatmap showing log₂ fold changes in LTR lineage abundance relative to corresponding diploid reference genomes. The following species names were shortened: *Ae. neglecta* subsp. *neglecta* is shown as *Ae. neglecta* (4x); *Ae. crassa* subsp. *crassa* (4x) as *Ae. crassa* (4x); *Ae. crassa* subsp. *crassa* (6x) as *Ae. crassa* (6x); and *Ae. neglecta* subsp. *recta* as *Ae. neglecta* (6x).

Despite similar overall TE content, variations in composition were observed among species. In particular, *Ae. crassa* subsp. *crassa* (4x), *Ae. vavilovii*, and *Ae. crassa* subsp. *crassa* (6x) had fewer *Gypsy* (38.00%, 38.90%, 39.71%) and *Copia* (13.97%, 14.90%, 14.72%), but more CACTA elements (16.01%, 14.32%, 15.29%) relative to other genomes. In addition, *Ae. crassa* subsp. *crassa* (4x) has the highest proportion of unclassified LTR elements (9.25%) (Supplemental Table 11). These results indicate lineage-specific differences in TE composition across *Aegilops* species.

### Comparative analysis of LTR clade abundance and dynamics across subgenomes

To assess LTR retrotransposon composition across *Aegilops* genomes, LTRs were classified into distinct clades using a domain-based HMM approach. Across all genomes, we identified four *Copia* (*Angela*, *SIRE*, *Ale*, *TAR*) and five *Gypsy* (*Retand*, *Athila*, *Tekay*, *CRM*, *Reina*) clades. For each clade, LTR counts were normalized per gigabase (Gb) to enable comparison across genomes and subgenomes (Supplemental Table 12), with polyploid subgenomes compared to their diploid references (Figures 5B-5G).

*Angela* was the most abundant *Copia* element, followed by *SIRE*, whereas *Ale* and *TAR* were less represented. Among *Gypsy* elements, *Retand* and *Athila* were predominant, with *Tekay* and *CRM* showing intermediate levels, and *Reina* remaining low. Overall, LTR composition was broadly conserved, though total abundance and specific clade proportions showed lineage- and subgenome-specific variations. M and D subgenomes showed the greatest variability, whereas U and S subgenomes closely resembled their diploid progenitors, with minor variation in C and N subgenomes (Figures 5B-5G and Supplemental Table 12).

Log₂ fold change analysis relative to diploid genomes revealed lineage-specific dynamics (Figure 5H). *Angela* consistently contracted in several M subgenomes including *Ae. crassa* subsp. *crassa* (4x and 6x) and *Ae. juvenalis*, while *SIRE* and *Ale* varied moderately, with *Ale* expanding in some D subgenomes. *TAR* displayed localized expansion in certain M subgenomes while remaining stable across other subgenomes. Among *Gypsy* lineages, *Retand* and *Athila* expanded in multiple M subgenomes, and *Tekay* contracted within the D lineage. *CRM* displayed contrasting patterns, with depletion in some U subgenomes such as *Ae. geniculata*, *Ae. biuncialis*, but expanded in several M and D subgenomes. In contrast, *Reina* consistently expanded across all D subgenomes (Figure 5H). Independent of species identity, fold-change patterns clustered subgenomes with similar LTR dynamics. While overall composition is conserved, individual LTR clades underwent subgenome-specific expansion and contraction.

### Subgenome-specific LTR dynamics within polyploid *Aegilops* species

To investigate LTR dynamics within polyploid genomes, we compared LTR clade abundance between subgenomes using log₂ fold-change. For tetraploids, subgenome pairs were compared (e.g., U vs M in *Ae. biuncialis*), while in hexaploids all pairwise comparisons were performed (e.g., D vs M, D vs S, M vs S in *Ae. vavilovii*). Overall, most clade-level differences were modest (log₂ fold-change < 0.5), indicating similar LTR composition within species (Figure 6; Supplemental Table 13).

**Figure 6.**
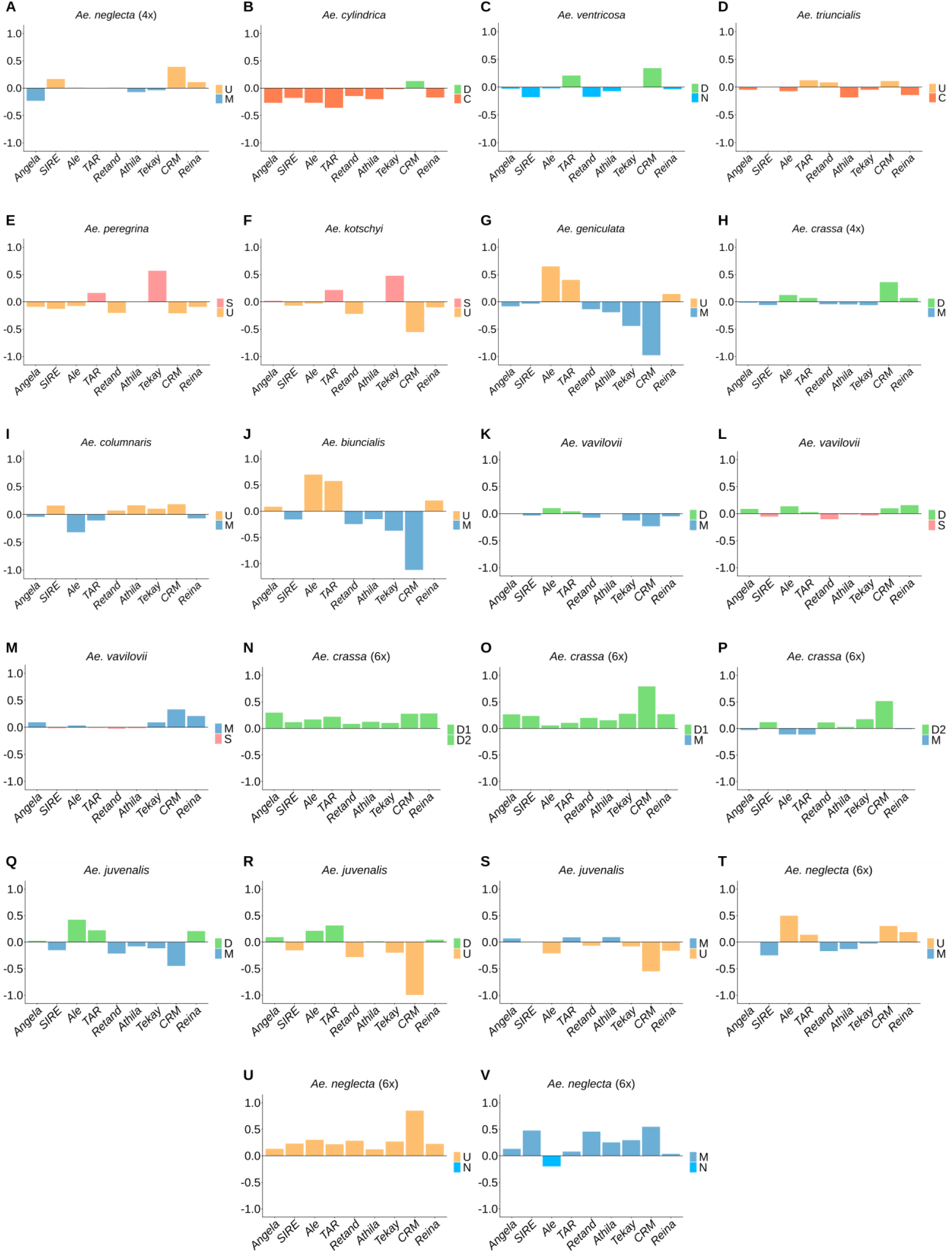
Subgenome-specific LTR dynamics within polyploid *Aegilops* species. **(A–V)** Log₂ fold changes in LTR lineage abundance between subgenomes within tetraploid and hexaploid species, highlighting subgenome-specific expansions and contractions of individual LTR lineages. For each panel, species names are at the top and subgenomes compared are specified in the legend. The following species names were shortened: *Ae. neglecta* subsp. *neglecta* is shown as *Ae. neglecta* (4x); *Ae. crassa* subsp. *crassa* (4x) as *Ae. crassa* (4x); *Ae. crassa* subsp. *crassa* (6x) as *Ae. crassa* (6x); and *Ae. neglecta* subsp. *recta* as *Ae. neglecta* (6x).

Despite this overall similarity, consistent subgenome-specific patterns were observed. In *Ae. geniculata* and *Ae. biuncialis*, the *Ale* and *TAR Copia* clades were enriched in the U subgenome, while the *CRM Gypsy* clade was consistently enriched in the M subgenome (Figures 6G and 6J), suggesting conserved dynamics. In *Ae. peregrina* and *Ae. kotschyi*, the *Tekay Gypsy* clade was enriched in the S subgenomes, with additional *CRM* expansion in the U subgenome of *Ae. kotschyi* (Figures 6E-6F). In tetraploids *Ae. neglecta* subsp. *neglecta*, *Ae. cylindrica*, *Ae. ventricosa*, *Ae. triuncialis*, *Ae. crassa* subsp. *crassa* (4x), and *Ae. columnaris*, minimal differences were observed for most clades, supporting overall homogenization (Figures 6A–6D and 6H–6I). However, localized expansions were still evident, including *TAR* in the C subgenome of *Ae. cylindrica* (Figure 6B), *Ale* in the M subgenome of *Ae. columnaris* (Figure 6I), and *CRM* enrichment in multiple U and D subgenomes (Figures 6A, 6C and 6H).

In hexaploids, subgenome-specific differences were limited but more pronounced in some species. In *Ae. juvenalis*, *CRM* showed clear expansion in the U subgenome relative to D and M (Figures 6Q–6S), while in *Ae. crassa* subsp. *crassa* (6x), its expansion was in D subgenomes relative to M (Figures 6N-6P). In *Ae. neglecta* subsp. *recta*, *CRM* was enriched in the U and M subgenomes compared with the N subgenome, whereas *Ale* expanded in the M subgenome relative to the U subgenome (Figures 6T–6V). In contrast, *Ae. vavilovii* showed minimal variation, indicating a highly homogenized LTR landscape (Figures 6K–6M). Overall, LTR composition is largely homogenized among subgenomes within species, likely reflecting shared nuclear environments after polyploidization. However, repeated expansion of specific lineages—particularly *Gypsy CRM*—across multiple species suggests ongoing or preferential activity in certain genomic contexts, highlighting its role in recent LTR dynamics in polyploid *Aegilops*.

### Insertion time dynamics of LTR retrotransposons across subgenomes

To investigate temporal dynamics of LTR retrotransposons, insertion times were estimated for Full-length elements in each subgenome and compared with their diploid references. Across all genomes, a consistent temporal separation between LTR superfamilies was observed. *Copia* elements showed predominantly recent insertions, peaking at ∼0–1 MYA, whereas *Gypsy* elements exhibited older insertion profiles, generally peaking at ∼0.5–1.5 MYA (Figure 7 and Supplemental Table 14).

**Figure 7.**
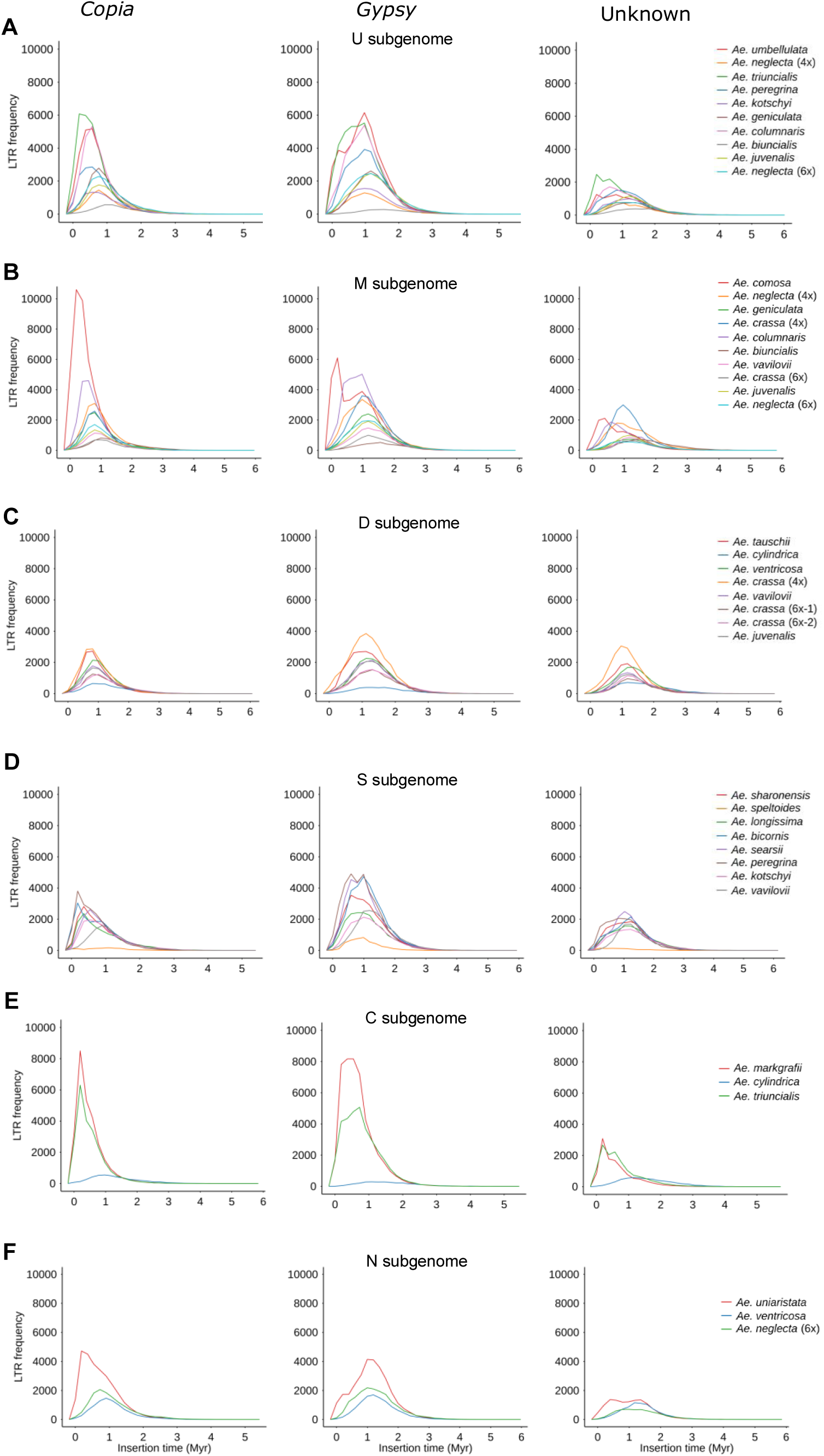
Insertion time dynamics of LTR retrotransposons across *Aegilops* subgenomes. **(A–F)** Insertion time distributions of LTR retrotransposons in **(A)** U, **(B)** M, **(C)** D, **(D)** S, **(E)** C, and **(F)** N subgenomes, including comparisons with the corresponding diploid reference genomes. The x-axes indicate insertion time in million years (Myr). The following species names were shortened: *Ae. neglecta* subsp. *neglecta* is shown as *Ae. neglecta* (4x); *Ae. crassa* subsp. *crassa* (4x) as *Ae. crassa* (4x); *Ae. crassa* subsp. *crassa* (6x) as *Ae. crassa* (6x); and *Ae. neglecta* subsp. *recta* as *Ae. neglecta* (6x). *Ae. crassa* (6x-1) and *Ae. crassa* (6x-2) denote the D1 and D2 subgenomes of *Ae. crassa* subsp. *crassa* (6x), respectively.

Subgenome-specific variation in LTR abundance was also evident. Several subgenomes, including the U and M subgenomes of *Ae. biuncialis*, the M subgenome of *Ae. crassa* subsp. *crassa* (6x), and the D and C subgenomes of *Ae. cylindrica*, exhibited lower numbers of full-length LTRs, indicating variation in intact LTR representation across subgenomes (Figures 7A–7C and 7E). Differences were also apparent at the ploidy level. Tetraploid subgenomes showed LTR insertion time distributions broadly similar to their diploid progenitors, whereas hexaploid genomes exhibited lower overall frequencies of full-length LTR elements and less pronounced peaks, indicating differences in the abundance and preservation of intact LTR retrotransposons across ploidy levels.

To place these patterns in an evolutionary context, insertion time distributions were compared with estimated divergence times between polyploid subgenomes and their corresponding diploid progenitors. Divergence time analysis indicated that subgenomes diverged more than 2 MYA (Figure 3), whereas many peak LTR insertions occurred within the last 0–1.5 MY (Figure 7 and Supplemental Table 14). This temporal separation indicates that most LTR activity postdates subgenome divergence and therefore represents lineage- and subgenome-specific transposition. Consistent with this, recent *Copia* activity (∼0–1 MYA) suggests ongoing amplification after polyploidization. Although *Gypsy* elements show slightly older insertion profiles, their activity also largely falls within the post-divergence period, suggesting independent amplification across lineages. Overall, these results indicate that LTR dynamics in *Aegilops* are driven primarily by both lineage-specific insertion histories and the differential retention or degradation of older elements.

## Discussion

We generated near-pseudomolecule-scale genome assemblies of four diploid, ten tetraploid, and four hexaploid species of *Aegilops*, which, together with the previously published ones, complete the production of assemblies for all 25 species of the genus. These assemblies substantially expand genomic resources for the genus, particularly for polyploid taxa, for which only *Ae. ventricosa* (Liu et al., 2025) was previously available. They provide a foundation for investigating genome evolution, polyploidization, structural variation, and traits relevant to wheat improvement.

Overall, assembly sizes are consistent with previous estimates while showing improved contiguity and completeness. Diploid assemblies closely match earlier reports (Avni et al., 2022; Li et al., 2022), with higher N50 and N90 values indicating improved continuity. Tetraploid genome sizes are concordant with cytogenetic estimates (Eilam et al., 2008; Eilam et al., 2007) and our flow cytometry data (Supplemental Table 15), with S^x^-containing species (e.g., *Ae. kotschyi*, *Ae. peregrina*) having larger genomes, whereas C-genome species (e.g., *Ae. cylindrica*, *Ae. triuncialis*) exhibit smaller genome sizes (Eilam et al., 2008; Eilam et al., 2007). The *Ae. ventricosa* assembly is comparable to that of Liu et al. (2025). The four hexaploid assemblies represent the first scaffold-level genomic resources for these species, with consistently high contiguity despite the challenges of assembling large, repetitive polyploid genomes. Comparative analyses further revealed a consistent pattern of genome downsizing across nearly all polyploid subgenomes relative to their diploid progenitors, indicating that genome contraction represents a pervasive feature of polyploid genome evolution in *Aegilops*. Similar patterns have been reported in other Triticeae polyploids and are generally interpreted as consequences of post-polyploid diploidization involving sequence elimination and genome restructuring (Eilam et al., 2008; Leitch and Bennett, 2004).

Building on these assemblies, protein-coding genes were annotated across all ploidy levels. High-confidence (HC) gene counts were generally lower than in previous studies. In diploids, HC gene numbers ranged from 21,865–27,783, compared with 31,183–40,222 reported in prior studies (Avni et al., 2022; Li et al., 2022; Yu et al., 2022). Similarly, tetraploid genomes had fewer HC genes than *Ae. ventricosa* (69,000) (Liu et al., 2025) and tetraploid wheat (65,012–66,559) (Avni et al., 2017; Maccaferri et al., 2019), although *Ae. triuncialis* (60,490) showed comparable values. In hexaploids, HC gene counts were also lower than the 106,913 genes reported for the Chinese Spring wheat genome (IWGSC, 2021). These differences could stem from methodological variation. Previous annotations incorporated RNA-Seq, Iso-Seq, and homology-based evidence, whereas our approach used a machine-learning model without transcriptomic support, but applied a stricter 95% similarity threshold (vs. 80–90% in earlier studies). Despite lower HC gene counts, BUSCO completeness (84.5%–98.9%) indicates that vast majority of the conserved genes were captured, supporting their use in comparative and evolutionary analyses.

Annotation of protein-coding genes in *Ae. crassa* subsp. *crassa* (4x) revealed higher gene content than in other tetraploid assemblies. Because elevated gene numbers can raise concerns about ploidy misclassification, nuclear DNA content was assessed by flow cytometry, confirming a genome size of 10.39 Gb, consistent with other tetraploids (8.83–12.01 Gb) (Supplemental Table 15). The increased gene number suggests lineage-specific gene retention and reduced post-polyploidization gene loss, a pattern commonly observed in polyploid plant genomes following whole-genome duplication (Cheng et al., 2018). Its deep phylogenetic divergence further suggests that extended evolutionary time contributed to the accumulation and retention of lineage-specific genes relative to other tetraploids.

Subgenome-level comparisons revealed that gene content evolution following polyploidization was highly heterogeneous. The M and D subgenomes displayed the greatest variability, including substantial gene loss in several lineages, consistent with the strongest genome downsizing observed in the M subgenomes. In comparison, the U and N subgenomes exhibited intermediate patterns characterized by moderate lineage-specific expansion and contraction, indicating a more limited but still detectable degree of post-polyploid remodeling. The S and C subgenomes remained comparatively conserved, retaining gene numbers similar to their diploid progenitors, consistent with genome size comparisons showing minimal deviation from diploid genome sizes in these lineages. Overall, these results indicate that genome downsizing following polyploidization is associated with widespread and uneven gene fractionation across subgenomes, reflecting coordinated but lineage-specific reductions in gene content.

The phylogenomic framework provides a comprehensive assessment of evolutionary relationships across all 25 recognized *Aegilops* species together with key *Triticum* genomes. The inferred relationships are largely consistent with established cytogenetic and taxonomic frameworks (Gill and Friebe, 2002; Kilian et al., 2011; Lilienfeld, 1951; Van Slageren, 1994) while also revealing lineage-specific complexities of genome evolution in the *Aegilops–Triticum* complex. Within the A clade, *T. urartu* and *T. monococcum* cluster with A subgenomes of polyploid wheats, consistent with their established roles as A-genome donors (Kilian et al., 2011; Li et al., 2022; Marcussen et al., 2014). The B-genome clade includes the B subgenomes of *T. aestivum* and *T. turgidum*, together with the S genome of *Ae. speltoides* and the G genome of *T. timopheevii*. The T genome of *Ae. mutica* also groups within this lineage, substantiating previous studies that suggested it as an early-diverging member of the B lineage. (Avni et al., 2022; Bernhardt et al., 2020; Glémin et al., 2019; Li et al., 2022)

The D genomes form a well-supported clade, with all D subgenomes clustering with *Ae. tauschii* and the D genome of hexaploid wheat, consistent with previous evidence for their origin (Chen et al., 2020; Lilienfeld and Kihara, 1934; Liu et al., 2025). Notably, divergence patterns suggest a more ancient origin of D subgenomes in polyploid *Aegilops* than previously proposed (Liu et al., 2025), with *Ae. crassa* subsp. *crassa* (4x) showing the earliest divergence, supporting its status as one of the oldest polyploid species (Dubcovsky and Dvorak, 1995; Kroupin et al., 2022). The “S” subgenomes of *Ae. peregrina* and *Ae. kotschyi* cluster with *Ae. longissima* (S^l^) and *Ae. sharonensis* (S^sh^), rather than *Ae. speltoides* (S), consistent with earlier studies (Badaeva et al., 2004; Friebe, 1996; Jewell, 1979; Jewell and Driscoll, 1983; Zhang et al., 1992; Zhao et al., 2016). Structural variation patterns further support *Ae. sharonensis* as the most likely progenitor (Supplemental Figures 2I and 2L and Supplemental Table 7). Similarly, the N, C, and U subgenomes group with their respective diploid donors (*Ae. uniaristata*, *Ae. markgrafii*, and *Ae. umbellulata*). For the N lineage, divergence estimates indicate a more ancient origin for *Ae. ventricosa* compared to previous reports (Liu et al., 2025) possibly reflecting an earlier split and subsequent independent evolution. Overall, these patterns support largely conserved relationships between polyploid subgenomes and their diploid progenitors.

The M genome displays the most complex and phylogenetically dispersed pattern. Only the M subgenomes of *Ae. geniculata* and *Ae. biuncialis* cluster with the diploid donor *Ae. comosa*, whereas others group within U- or D-genome clades. This pattern contrasts with the expected donor–subgenome relationships and suggests a more complex evolutionary history than previously assumed. Most M subgenomes show deep divergence (>4 MYA), placing them among the oldest polyploid-derived subgenomes in *Aegilops*. Notably, the M subgenome of *Ae. crassa* subsp. *crassa* (4x; DM) shows the earliest divergence, consistent with earlier cytogenetic studies identifying this species as one of the most ancient polyploids (Dubcovsky and Dvorak, 1995; Kroupin et al., 2022). This phylogenetic displacement, together with deep divergence, suggests post-polyploidization extensive genomic remodeling, likely driven by structural variation and homoeologous exchanges during prolonged coexistence with D or U genomes (Feldman and Levy, 2012; Zhang et al., 2023).

Comparative analyses of the assemblies revealed that extensive structural variation is a pervasive feature of polyploid *Aegilops* genomes and a major driver of post-polyploidization evolution. While many subgenome–progenitor comparisons retain large syntenic regions, widespread inversions, translocations, duplications, insertions, and deletions indicate genome reorganization following polyploid formation, consistent with observations in wheat and other *Aegilops* polyploids (Badaeva et al., 2004; Badaeva et al., 2021; Friebe, 1996).

Large inversions, though less frequent, produce major structural changes, while abundant translocations reflect homoeologous pairing and exchanges documented cytogenetically (Badaeva et al., 2004; Friebe, 1996; Lilienfeld, 1951), indicating that interchromosomal rearrangements have been a persistent feature of their evolutionary history. Duplications, the most prevalent SV type, often involve large segments and likely contribute to gene copy number variation, dosage effects, and functional diversification, all recognized sources of evolutionary novelty in polyploid genomes (Flagel and Wendel, 2009).

Patterns of structural variation closely mirror phylogenetic relationships. Subgenomes phylogenetically displaced from their diploid progenitors—particularly M subgenomes—show reduced synteny and few detectable SVs in direct comparisons, whereas comparisons with references of co-resident subgenomes reveal higher synteny and more SVs, suggesting that prolonged coexistence of divergent subgenomes facilitates interchromosomal rearrangements and homoeologous exchanges, widening divergence from diploid progenitors.

Comparisons between subgenomes within individual polyploid species further supported extensive post-polyploidization genome differentiation. While most subgenome pairs exhibited relatively low levels of synteny, indicating substantial structural divergence, *Ae. triuncialis, Ae. columnaris* and tetraploid *Ae. crassa* retained greater collinearity despite harboring large numbers of translocations and duplications. These findings suggest that the extent of subgenome restructuring varies among polyploid species and that prolonged coexistence within the same genome can lead to distinct evolutionary trajectories for different genomic combinations.

Considering the limited availability of comprehensive pan-genome resources for *Aegilops* species, it is not possible to unambiguously distinguish structural variants that are unique to polyploidization from those already present within natural diploid populations. However, additional comparisons among multiple diploid *Aegilops* genomes from the same species indicate that substantial structural variation is already present within diploid gene pools. This suggests that at least a portion of the SV observed in polyploid–progenitor comparisons may reflect pre-existing variation within ancestral diploid populations, in addition to structural changes accumulated following polyploidization.

Genome-wide TE annotation across 18 *Aegilops* species confirms that TEs constitute a major fraction of these genomes (85–88%), dominated by LTR retrotransposons (*Gypsy* and *Copia*) and *CACTA* DNA transposons, consistent with other Triticeae species (Vicient and Casacuberta, 2017;

Wicker et al., 2018). While overall TE composition is broadly conserved, lineage-specific variations—such as reduced *Gypsy* and *Copia* content and elevated *CACTA* abundance in several polyploids—highlight differential retention and amplification, reflecting the dynamic behavior of TEs following polyploidization (Fedoroff, 2000; Parisod et al., 2010).

Comparative analysis of LTR retrotransposon clades across *Aegilops* genomes revealed TE landscape dominated by a limited number of highly abundant lineages. Across all genomes, *Angela* represented the most abundant *Copia* element, whereas within the *Gypsy* superfamily, *Retand* and *Athila* were consistently predominant, indicating that these clades constitute the major components of LTR retrotransposon diversity in *Aegilops*. Comparative LTR analyses across subgenomes relative to their diploid progenitors reveal that post-polyploidization TE dynamics are highly lineage- and subgenome-specific, rather than following a uniform trajectory. Instead, distinct LTR clades exhibit contrasting patterns depending on genomic context. For example, the *Copia* element *Angela* shows consistent contraction across most M subgenomes, whereas *TAR* displays localized expansions in specific M subgenomes, indicating heterogeneous *Copia* dynamics within the same subgenomic background. In contrast, the *Gypsy* lineage *Reina* shows a consistent expansion across all D subgenomes, suggesting a shared D-genome-specific propensity for this TE family. Similarly, *CRM* exhibits highly contrasting behavior across genomes, with contraction in several U subgenomes (e.g., *Ae. biuncialis* and *Ae. geniculata*), but expansion in selected M and D subgenomes. These lineage-specific LTR dynamics were broadly consistent with patterns of genome size evolution across subgenomes. Subgenomes exhibiting stronger genome downsizing, particularly the M subgenomes, generally also showed greater shifts in LTR composition and abundance, suggesting that TE turnover contributed, at least in part, to genome contraction following polyploidization. In contrast, the S subgenomes remained comparatively stable in LTR composition and abundance, closely resembling their diploid progenitors, consistent with their overall conservation in genome size and gene content. Together, these findings indicate that different subgenomes followed distinct evolutionary trajectories after polyploidization, with TE dynamics contributing differentially to genome remodeling and subgenome divergence in *Aegilops*.

Subgenome-level analyses within polyploids further indicate that while most LTR clades are homogenized, certain families display localized expansions or contractions. Notably, the *Gypsy CRM* consistently exhibits subgenome-specific expansion, suggesting preferential activity or retention. Temporal analyses show *Copia* elements driving recent genome evolution, whereas *Gypsy* elements largely reflect older amplification events, likely influenced by transposition activity, epigenetic regulation, and element removal via recombination or deletion (Bennetzen and Wang, 2014; Fedoroff, 2000; Lisch, 2013; Pulido and Casacuberta, 2023). Overall, these findings underscore the central role of TEs in shaping *Aegilops* genome structure and their contribution to subgenome differentiation and polyploid genome evolution.

In conclusion, we provide a comprehensive genomic framework for 18 *Aegilops* species, integrating high-quality assemblies, gene annotation, phylogenetic, and structural variation analyses. Polyploid subgenomes show substantial divergence from their diploid progenitors, driven by extensive interchromosomal remodeling. LTR analyses reveal lineage- and subgenome-specific expansions and contractions, highlighting the dynamic role of TEs upon polyploidization. The multi-faceted analysis provides insights into polyploid *Aegilops* genome evolution and can be capitalized upon to inform wheat improvement through novel alleles, subgenome variation, and structural diversity.

## Methods

### Plant materials

One accession each of 18 *Aegilops* species (Supplemental Table 1) were grown in root trainers in a growth cabinet with a 16 h light and 8 h dark photoperiod regime, and temperature settings of 24°C during the day and 18°C at night. Seedlings were vernalized at the 4-5 leaf stage for ten weeks in a 4°C growth chamber with a 12 h photoperiod. After vernalization, seedlings were transplanted and returned to the original growth chamber. At heading, plants were bagged to prevent cross-pollination and seeds were harvested at maturity. Single-seed descents were grown for 3-5 generations to increase homozygosity prior to genome sequencing initiation.

### Isolation of high-molecular-weight genomic DNA

Ten to 58 seedlings from each of the 18 single-seed descent accessions (Supplemental Table 1) were grown in root trainers as described above. At the 2-4 leaf stage, plants were maintained in darkness for 2-3 days, after which tissue collection was performed by weighing the tissue prior to flash-freezing and storage in liquid nitrogen. Seedlings were allowed to regrow and the sampling process was repeated until approximately 15 g was collected per sample.

High-molecular-weight genomic DNA (HMW gDNA) was extracted using a modified nuclei isolation protocol (Zhang et al., 2012) where the nuclei isolation buffer (NIB) was supplemented immediately before use with spermine trihydrochloride and spermidine tetrahydrochloride to final concentrations of 1 mM each. Briefly, the tissue was ground to a fine powder in liquid nitrogen using a mortar and pestle and homogenized in the supplemented NIB. The homogenate was filtered and divided into four 50 mL Falcon tubes. Nuclei were pelleted by centrifugation at 2,786 *g* for 20 minutes at 4°C. Pellets were gently resuspended in 200 µL ice-cold NIB using a paintbrush and combined into a single 50 mL Falcon tube. The nuclei were washed four additional times with 40 mL supplemented NIB. After the final wash, nuclei were resuspended in 20 mL of 1× homogenization buffer (HB) containing 1% (w/v) polyvinylpyrrolidone (PVP40), followed by centrifugation. The final pellet was gently resuspended in 3 mL of 1× HB + PVP using minimal pipetting and a wide-bore tip.

The nuclei suspension was divided into six 15-mL Falcon tubes (∼500 µL per tube), and nuclei were pelleted by centrifugation at 2,450 *g* for 10 minutes at 4°C. Each pellet was resuspended in 3.6 mL lysis buffer (100 mM Tris-HCl pH 9.0, 25 mM Na₂EDTA, 138 mM NaCl, 1% PVP40, 0.3 mg/mL proteinase K, 1% sodium lauryl sarcosine) and incubated at 50°C with gentle agitation for 75–90 minutes, with manual inversion every 10 minutes. Following lysis, 1.4 mL 5M NaCl was added (final concentration 1.5 M), and samples were gently mixed and centrifuged at 2,755 *g* for 15 minutes at 4°C to pellet the debris. The gDNA-containing supernatants were transferred to new tubes, and the DNA was precipitated by adding 2.5 volumes cold absolute ethanol (∼11 mL). Tubes were gently rocked for 1-2 min until DNA strands were visible, followed by mixing on a HulaMixer (ThermoFisher Scientific, Waltham, MA, USA) for an additional 10 minutes. The DNA was pelleted by centrifugation at 4,260 *g* for 15 minutes at 4°C. The DNA pellets were washed with 10 mL cold 70% ethanol, mixed again on the HulaMixer for ten minutes, centrifuged, and air-dried horizontally in a laminar flow hood for 30 minutes. Any residual ethanol was carefully removed with a long-stem swab. A total of 200 µL “low-TE” buffer (10 mM Tris, 0.1 mM EDTA, pH 8.0) was added to each tube, mixed by gently flicking the tubes and stored at 4°C for 2–4 days to allow complete resuspension of the HMW gDNA before quantification using the Quant-iT™ PicoGreen® dsDNA assay kit (ThermoFisher Scientific, Burlington, ON, Canada).

DNA quality was assessed by pulsed-field gel electrophoresis (PFGE). DNA samples were loaded on a 1% UltraPure™ agarose gel (ThermoFisher Scientific) in 0.5× TBE buffer and resolved on a CHEF DR-II electrophoresis system (Bio-Rad, Mississauga, Ontario, Canada) at 6 V/cm with a pulse time from 5 to 15 seconds over 16 hours at 12.5°C. Lambda PFG Ladder and Midrange PFG Marker (New England Biolabs, Ipswich, MA, USA) were used as molecular size standards.

### Sequencing

Multiple library types were constructed depending on the species: (1) Oxford Nanopore Technologies (ONT) long-read, (2) PacBio SMRTbell, (3) Illumina short-read, (4) Hi-C and (5) Omni-C. Libraries were sequenced on the corresponding platforms to achieve the following coverages: 21–63x for ONT, 3–6x for PacBio HiFi, 10–23× for Illumina short reads, 10–40x for Hi-C, and 9–23x for Omni-C (Supplemental Table 2).

### Oxford Nanopore Technologies (ONT)

ONT 1D ligation libraries were prepared using the LSK109 kit, following the manufacturer’s protocol with minor modifications to maximize retention of ultra-long DNA fragments. The HMW gDNA was size-selected using a BluePippin system (Sage Science, MA, USA) with the high-pass protocol to enrich for fragments larger than 30 kb. Following size selection, magnetic bead purification was used to clean and concentrate the DNA. The concentrations were determined using a Qubit 2.0 Fluorometer (ThermoFisher Scientific, Burlington, ON, Canada), and integrity was assessed using a TapeStation 2200 system (Agilent Technologies, Mississauga, ON, Canada). The samples were stored at 4°C until library preparation. For each library, 1.2–1.5 µg of end-repaired, size-selected HMW gDNA was used. Sequencing was performed on a PromethION (Oxford Nanopore Technologies, Oxford, UK) using R9 flow cells with high accuracy base calling.

### PacBio HiFi

For PacBio sequencing, HMW gDNA was sent to the Centre d’expertise et de services Génome Québec (Montréal, QC, Canada). PacBio gDNA libraries were prepared following the Pacific Biosciences SMRTbell® prep kit 3.0 protocol for whole genome and metagenome library construction. The HMW gDNA was sheared using the Diagenode Megaruptor 3 instrument (Diagenode Inc., Denville, NJ, USA). DNA damage repair, end repair, and SMRTbell adapter ligation steps were performed according to the manufacturer’s instructions with SMRTbell Template Prep Kit 3.0 reagents (Pacific Biosciences). Libraries were size-selected using a diluted AMPure PB bead protocol. Sequencing primers (version 3.2) were annealed to the libraries, and the Sequel II 2.2 polymerase was bound to the templates. Libraries were cleaned using SMRTbell cleanup beads following the SMRTlink v11 calculator protocol. Sequencing was performed on a PacBio Sequel II instrument at a loading concentration of 80 pM using the adaptive loading protocol, Sequel II Sequencing Kit 2.0, SMRT Cell 8M, with a 2-hour pre-extension time and a 30-hour movie collection time.

### Illumina short reads

HMW gDNA at 50ng/µl was sent to the Centre d’expertise et de services Génome Québec (Montréal, QC, Canada). Libraries were prepared from 700 ng gDNA using the TruSeq DNA PCR-Free Library Preparation Kit (Lucigen, Wisconson, USA), following the manufacturer’s recommended protocol. Library quantification was performed using the KAPA Library Quantification Kit—Complete (Universal) (Kapa Biosystems). The average fragment size of each library was determined using the Agilent 5300 Fragment Analyzer. Libraries were normalized, pooled, and denatured in 0.02 N NaOH, then neutralized with HT1 buffer. The pooled libraries were loaded at 400 pM onto an Illumina NovaSeq 6000 S4 flow cell using the Xp workflow according to the manufacturer’s instructions. Sequencing was conducted in paired-end mode (2× 150 bp cycles). A 1% PhiX control library served as internal quality control. Base calling and demultiplexing were performed using BCL Convert version 4.2.4, generating high-quality FASTQ files for downstream analysis.

### Hi-C

Hi-C libraries were prepared following the protocol described by Padmarasu et al. (2019). Fresh young leaf tissue was crosslinked with formaldehyde, and chromatin was digested using the restriction enzyme *Dpn*II. The restricted DNA was blunt-end repaired and proximity ligation was performed, incorporating biotinylated nucleotides to enrich for ligation junctions. The resulting DNA was purified and used for library construction following the Illumina DNA Prep protocol (https://support-docs.illumina.com/LP/IlluminaDNAPrep/Content/LP/FrontPages/IlluminaDNAPrep.htm). Libraries were sent to the Centre d’expertise et de services Génome Québec (Montréal, QC, Canada) for sequencing.

At the sequencing center, libraries were indexed using TruSeq LT single indices (Illumina), PCR amplified and quantified using the KAPA Library Quantification Kit—Complete (Universal) (Kapa Biosystems), and average fragment size was confirmed on an Agilent 5300 Fragment Analyzer. Libraries were normalized, pooled (3–4 libraries per pool), denatured in 0.02 N NaOH, and neutralized using a pre-load buffer. The pooled libraries were loaded at 200 pM onto an Illumina NovaSeq 6000 S4 flow cells. Sequence and processing were as described for Illumina short reads above.

### Omni-C

Young leaf tissue was harvested from seedlings at the 2–4 leaf stage, following a dark treatment of approximately 72 hours. A total of 300 mg of fresh leaf tissue was collected and immediately frozen in liquid nitrogen. Omni-C libraries were prepared using the Dovetail® Omni-C® Kit (Cantata Bio, Scotts Valley, CA, USA), following the manufacturer’s user guide with plant-specific modifications. To obtain a good fragment size distribution, the Nuclease Enzyme Mix (Nucl C) was further diluted 2-4 folds relative to the standard Nucl C working concentration. Final Omni-C libraries were quantified using the Qubit™ dsDNA High Sensitivity Assay Kit (ThermoFisher Scientific), and fragment size distributions were assessed using a TapeStation 4200 system with High Sensitivity D5000 ScreenTape (Agilent Technologies). Libraries were sequenced at the Centre d’expertise et de services Génome Québec (Montréal, QC, Canada) using the same Illumina protocol described for Hi-C libraries above, including indexing, PCR amplification, quantification, and fragment-size assessment. Sequencing was performed on an Illumina NovaSeq 6000 S4 flow cell in 2 × 150 bp paired-end mode across two lanes: Lane 1 included libraries for *Ae. geniculata*, *Ae. juvenalis*, and *Ae. vavilovii*; Lane 2 included libraries for *Ae. crassa* subsp. *crassa* (6x), *Ae. vavilovii*, and *Ae. neglecta* subsp. *recta*. A 1% PhiX library was used as sequencing control, and raw data processing was performed as described for Hi-C libraries.

### Genome assembly

ONT long reads were assembled using the SMARTdenovo v1.0.0 (Liu et al., 2021) for all accessions except for *Ae. crassa* ssp. *crassa* (4x), *Ae. columnaris*, and *Ae. triuncialis*, which were assembled with hifiasm v0.19.8 (Cheng et al., 2021) due to suboptimal assemblies with SMARTdenovo. Initial draft assemblies were first polished with PacBio long reads using Inspector (Chen et al., 2021), followed by a second round of polishing with Illumina short reads using HyPo (Kundu et al., 2019). Hi-C data were used to scaffold the polished assemblies with YaHS (Zhou et al., 2022). Upon Hi-C scaffolding, assemblies of *Ae. geniculata*, *Ae. vavilovii*, *Ae. crassa* subsp. *crassa* (6x), *Ae. juvenalis*, and *Ae. neglecta* subsp. *recta* were more fragmented than the other 13; hence, Omni-C data were generated for these species, and scaffolding was performed with the HapHiC tool (Zeng et al., 2024). Assembly completeness and quality were assessed using BUSCO v5.2.2 (Manni et al., 2021), with the embryophyta_odb10 database comprising 1,614 conserved single-copy orthologs.

### Gene annotation

*Ab initio* gene prediction of all *Aegilops* genome assemblies was performed using Helixer v0.3.5, a deep learning-based tool for gene identification, using the land_plant lineage model with a peak threshold of 0.9 (Stiehler et al., 2020). Predicted gene models were subsequently classified into high-confidence (HC) and low-confidence (LC) categories based on sequence homology to curated protein databases. All predicted gene models were compared against UniPoa (Poaceae database of annotated proteins from the UniProt database), the UniMag database (validated Magnoliophyta proteins from SwissProt) (Boeckmann et al., 2003), and PTREP (a database of hypothetical proteins derived from the non-redundant set of transposable elements in TREP [Triticeae Repeat Sequence]) (Wicker et al., 2002). Sequence homology to each database was determined using BLASTP (https://blast.ncbi.nlm.nih.gov/Blast.cgi) with e-value < 10e^−5^. Classification into HC and LC gene categories was based on the best hit obtained using the following criteria: (i) alignments with at least 95% overlap between query and subject sequences for UniPoa and UniMag, and (ii) alignments with at least 95% query coverage for PTREP. Gene models were classified as HC if they had a significant hit in UniPoa and/or UniMag and were not present in PTREP. Gene models that did not meet these criteria were designated as LC.

### Orthogroup inference and pangenome analysis

Protein sequences derived from the predicted gene models of all 25 *Aegilops* species were used for orthogroup inference using OrthoFinder v3.1.0 (Emms and Kelly, 2019). Orthogroups were defined as sets of genes descended from a single gene in the last common ancestor. Based on their distribution across species, orthogroups were classified into three categories: (i) core orthogroups, present in all 25 species; (ii) dispensable orthogroups, present in two or more but not all species; and (iii) species-specific orthogroups, present in only one species. Corresponding genes within each category were assigned based on their inclusion in these orthogroups.

### Phylogenetic tree construction and divergence time estimation

Phylogenetic relationships were inferred from single-copy orthologs identified using OrthoFinder v3.1.0 (Emms and Kelly, 2019). For polyploid species, subgenomes were separated and analyzed as independent entries. The dataset comprised 36 *Aegilops* genomes and subgenomes generated in this study, along with 27 previously published *Aegilops* and *Triticum* genomes/subgenomes. The outgroup species *Hordeum vulgare* (Beier et al., 2017), *Thinopyrum elongatum* (Wang et al., 2020), and *Brachypodium distachyon* (Vogel et al., 2010), were included to root the phylogenetic tree. Reference genome assemblies for the previously published *Triticum/Aegilops* species were *T. urartu* (Ling et al., 2018), *T. monococcum* TA10622 and TA299 (Ahmed et al., 2023), A and B subgenomes of *T. turgidum* (Zhu et al., 2019), A and G subgenomes of *T. timopheevii* (Grewal et al., 2024), A, B and D subgenomes of *T. aestivum* (IWGSC, 2021), *Ae. tauschii* (Wang et al., 2021), *Ae. speltoides* (Yang et al., 2023), *Ae. sharonensis*, *Ae. longissima, Ae. speltoides* (Avni et al., 2022), *Ae. speltoides*, *Ae. bicornis*, *Ae. longissima*, *Ae. searsii*, *Ae. sharonensis* (Li et al., 2022), *Ae. mutica* (Grewal et al., 2025), *Ae. comosa* (Li et al., 2024), *Ae. umbellulata* (Abrouk et al., 2023), *Ae. comosa*, *Ae. umbellulata*, *Ae. markgrafii* and *Ae. uniaristata* (Khadka et al. 2026, unpublished). Because of the large size and diversity of this 66 genome dataset, no strict set of universal single-copy orthologs was initially identified. To address this, orthologous groups were defined as follows: single-copy genes must be present in at least 80% of the genomes, while allowing the remaining 20% of genomes to contain multiple copies. In the latter cases, only the longest isoform was retained for analysis. This approach yielded a final set of 773 near single-copy orthologs. Protein sequences from each orthologous group were aligned using MAFFT (Katoh and Standley, 2013), and the resulting alignments were back-translated to coding sequences (CDSs) with PAL2NAL (Suyama et al., 2006). Finally, conserved regions of each CDS alignment were extracted using Gblocks v0.91b (Castresana, 2000).

For phylogenetic inference, the filtered CDS alignments were concatenated into a single supermatrix using PhyKIT (https://jlsteenwyk.com/PhyKIT/usage/index.html#create-concatenation-matrix). Phylogenetic trees were constructed using the maximum-likelihood method implemented in RAxML-NG v1.2.2 (Kozlov et al., 2019) with the GTR+I+GAMMA model and 1000 bootstrap replicates.

Divergence times were estimated using a relaxed molecular clock implemented in MCMCTree from the PAML v4.10.9 package (Dos Reis and Yang, 2019), using the correlated rate model (clock = 3) and the HKY85 substitution model. Because evolutionary rates differ across codon positions, the concatenated supermatrix was partitioned by codon position, treating the first, second, and third positions as independent partitions. The MCMC analysis was run for 10,000,000 iterations following a burn-in of 3,000,000 iterations. Convergence was assessed by verifying that the effective sample size (ESS) for all parameters exceeded 200. Date calibrations were applied to major nodes based on previously published estimates: (i) the divergence between *Brachypodium distachyon* and the *Triticum*/*Aegilops* lineage (mean 44.4 ± 3.53 MYA) (Marcussen et al., 2014), (ii) the divergence between *Hordeum vulgare* (barley) and *Triticum*/*Aegilops* (∼13 MYA) (Gaut, 2002), and (iii) the divergence between the A- and B-genome lineages of wheat and *Aegilops* (∼7 MYA) (Marcussen et al., 2014). For each dataset, two independent MCMC runs were performed to confirm consistency and convergence of the posterior estimates. Tree topologies were visualized using FigTree v.1.4.4 (https://github.com/rambaut/figtree).

### Identification of structural variation

Previously generated diploid *Aegilops* genome assemblies for *Ae. comosa*, *Ae. umbellulata*, *Ae. markgrafii*, *Ae. uniaristata* (Khadka et al. 2026, unpublished), *Ae. speltoides* (Avni et al., 2022) and *Ae*. *tauschii* (Wang et al., 2021) were used as reference genomes for structural variation (SV) analysis. For each polyploid genome, the corresponding diploid progenitor genomes served as references. Subgenomes within the polyploid assemblies were assigned based on gene collinearity with their respective diploid progenitor genomes. Coding sequences from the diploid progenitor assemblies were mapped to the polyploid assemblies using GMAP (Wu and Watanabe, 2005) with minimum identity and coverage thresholds of 70%. Gene collinearity information was processed with a custom Perl script to assign scaffolds to their corresponding subgenomes. The seven largest scaffolds from each subgenome were selected for SV analyses. Scaffolds were renamed based on synteny with the assigned reference genome and standardized to match the chromosome naming of the diploid progenitor species. Pairwise whole-genome alignments between subgenomes and their respective diploid progenitors were performed using MUMmer v3.23 with the parameters --maxmatch -l 500 (Marçais et al., 2018). The alignment output was processed with the command line show-coords -THrd to generate coordinate files, which were then used as input for SV detection using the SyRI v1.7.0 (Goel et al., 2019) pipeline with default settings. All SVs > 50 bp were tallied. Visualization of SV plots was performed using the plotsr tool (Goel and Schneeberger, 2022).

### Transposable element identification and LTR analysis

Transposable elements (TEs) were annotated using the Extensive *de novo* TE Annotator (EDTA) v2.2.2 pipeline, which performs genome-wide *de novo* identification and classification of repetitive sequences (Ou et al., 2019). LTR retrotransposons identified by EDTA were further classified into lineages using TEsorter v1.4.7 (Zhang et al., 2022) with the rexdb-plant database (Neumann et al., 2019) and the 70-30-80 rule, assigning elements to specific superfamilies and clades based on conserved protein domains detected through HMM-based searches. Insertion times of intact LTR retrotransposons were estimated from the divergence between their 5′ and 3′ LTR sequences, as implemented in EDTA, assuming a substitution rate of 1.3 × 10⁻⁸ substitutions per site per year, following estimates from rice (Ou et al., 2019).

## Data availability

All 18 genome assemblies generated in this study have been deposited at the National Center for Biotechnology Information under BioProject PRJNA1449007. Accession numbers are as follows: *Ae. sharonensis* (xxx), *Ae. longissima* (xxx), *Ae. bicornis* (xxx), *Ae. searsii* (JBXSOC000000000), *Ae. neglecta* subsp. *neglecta* (JBXSOD000000000), *Ae. cylindrica* (JBXSOE000000000), *Ae. ventricosa* (JBXSQR000000000), *Ae. triuncialis* (JBXSQS000000000), *Ae. peregrina* (JBXSQT000000000), *Ae. kotschyi* (JBXSQU000000000), *Ae. geniculata* (JBXSQV000000000), *Ae. crassa* subsp. *crassa* (4x) (JBXSQW000000000), *Ae. columnaris* (JBXSQX000000000), *Ae. biuncialis* (JBXSQY000000000), *Ae. vavilovii* (JBXUDD000000000), *Ae. crassa* subsp. *crassa* (6x) (JBXUOA000000000), *Ae. juvenalis* (JBXUOB000000000), and *Ae. neglecta* subsp. *recta* (JBXUOC000000000). All data supporting the findings of this study are available within the article and its Supplementary Information.

## Author contributions

H.S. performed data analyses and interpretation, generated figures, and drafted the manuscript. T.E., M.L.L., and J.E. performed extraction, library construction, and data management. C.Z. performed data management. F.M.Y., C.J.P., and S.C. performed project management, study design, data generation and analysis, and manuscript editing. All co-authors read and approved the final manuscript.

## Acknowledgements

The authors gratefully acknowledge Genome Prairie for administrative support and thank Raelene Regier for coordination and assistance. We also thank Dr. Sara Martin and Tracey James for assistance with flow cytometry.

## Supplemental information

**Supplemental Figure 1.**
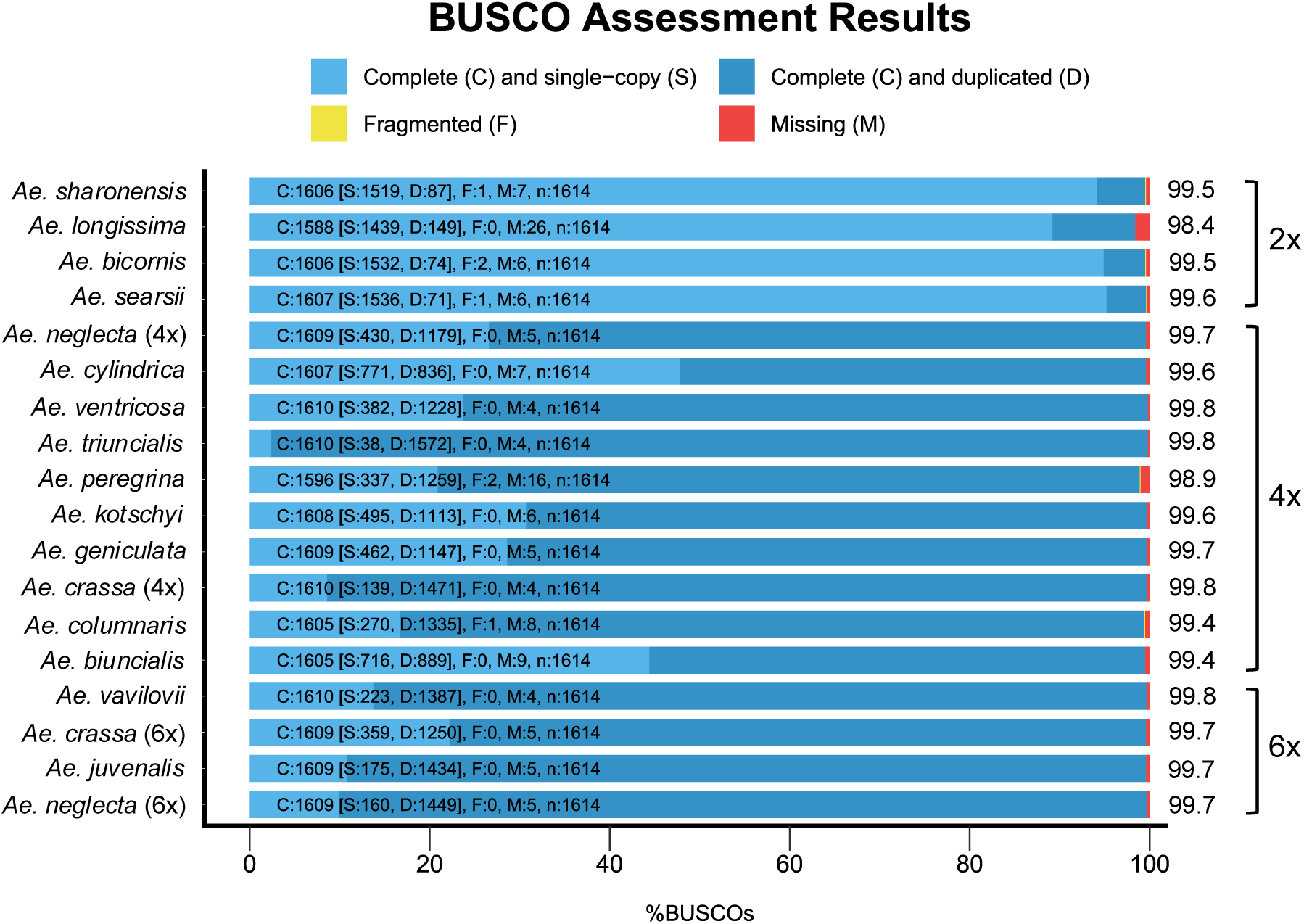
BUSCO assessment of genome assembly completeness in *Aegilops* genomes. Genome completeness was evaluated using the *embryophyta_odb10* database (n = 1,614 core plant genes). Each bar represents the proportion of BUSCOs classified as complete (C)[single-copy (S) or duplicated (D)], fragmented (F), or missing (M). The numbers inside each segment of the bars indicate the count of BUSCO genes in each category, while the numbers on the right of each bar show the corresponding BUSCO completeness score as a percentage. Genome ploidy, indicated on the right as 2x, 4x, and 6x, corresponds to diploid, tetraploid, and hexaploid assemblies, respectively. The following species names were shortened: *Ae. neglecta* subsp. *neglecta* is shown as *Ae. neglecta* (4x); *Ae. crassa* subsp. *crassa* (4x) as *Ae. crassa* (4x); *Ae. crassa* subsp. *crassa* (6x) as *Ae. crassa* (6x); and *Ae. neglecta* subsp. *recta* as *Ae. neglecta* (6x).

**Supplemental Figure 2.**
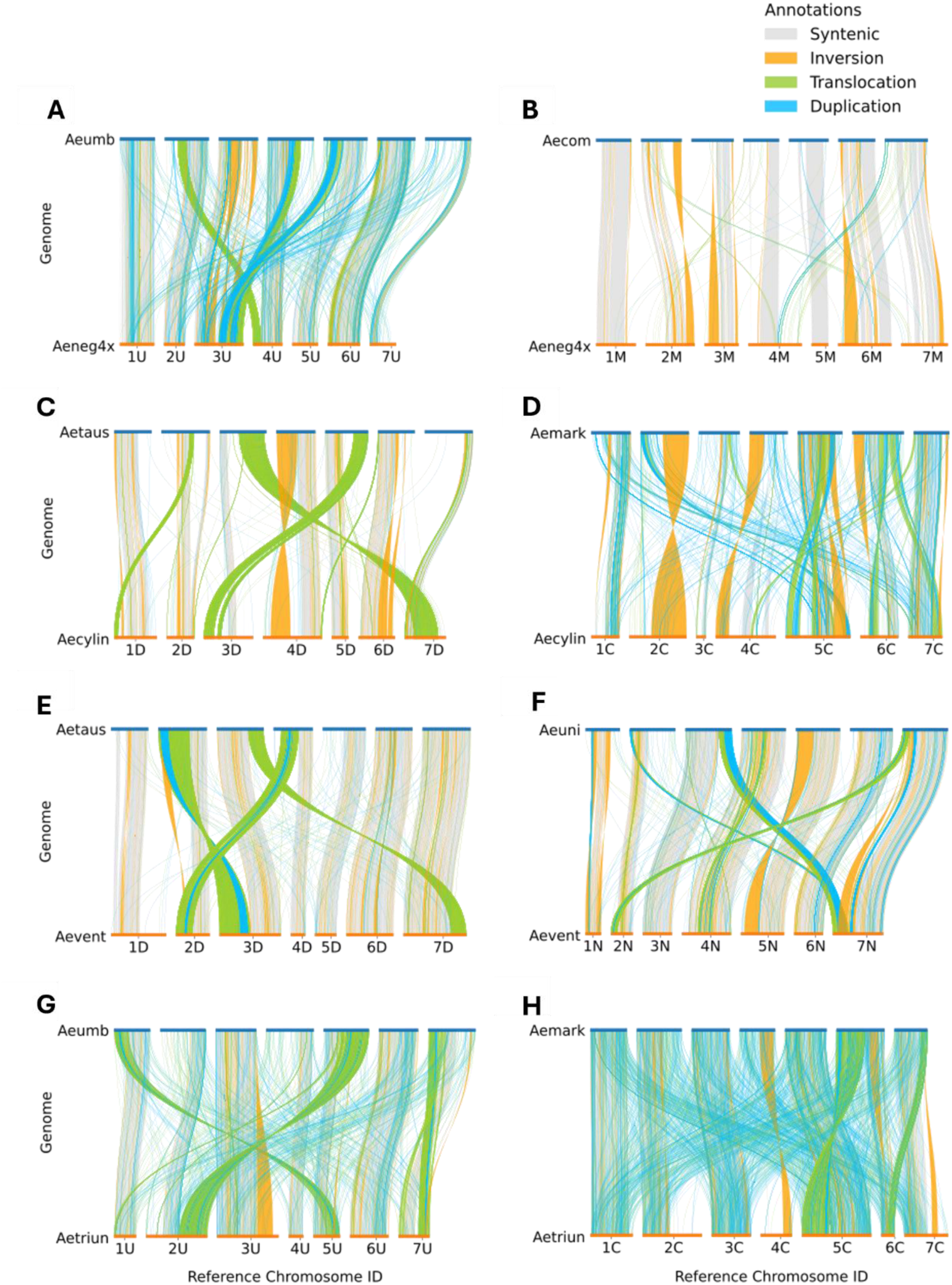
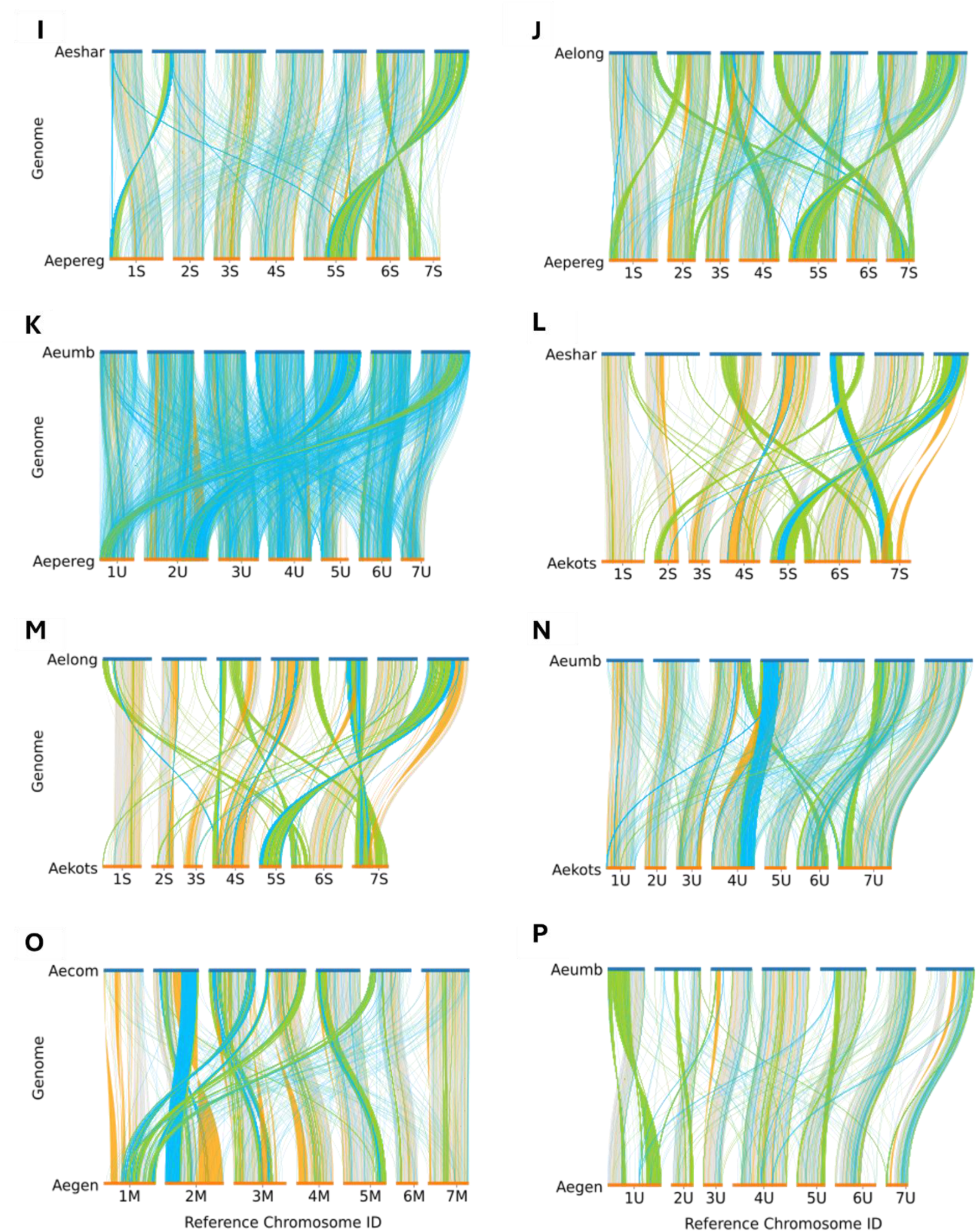
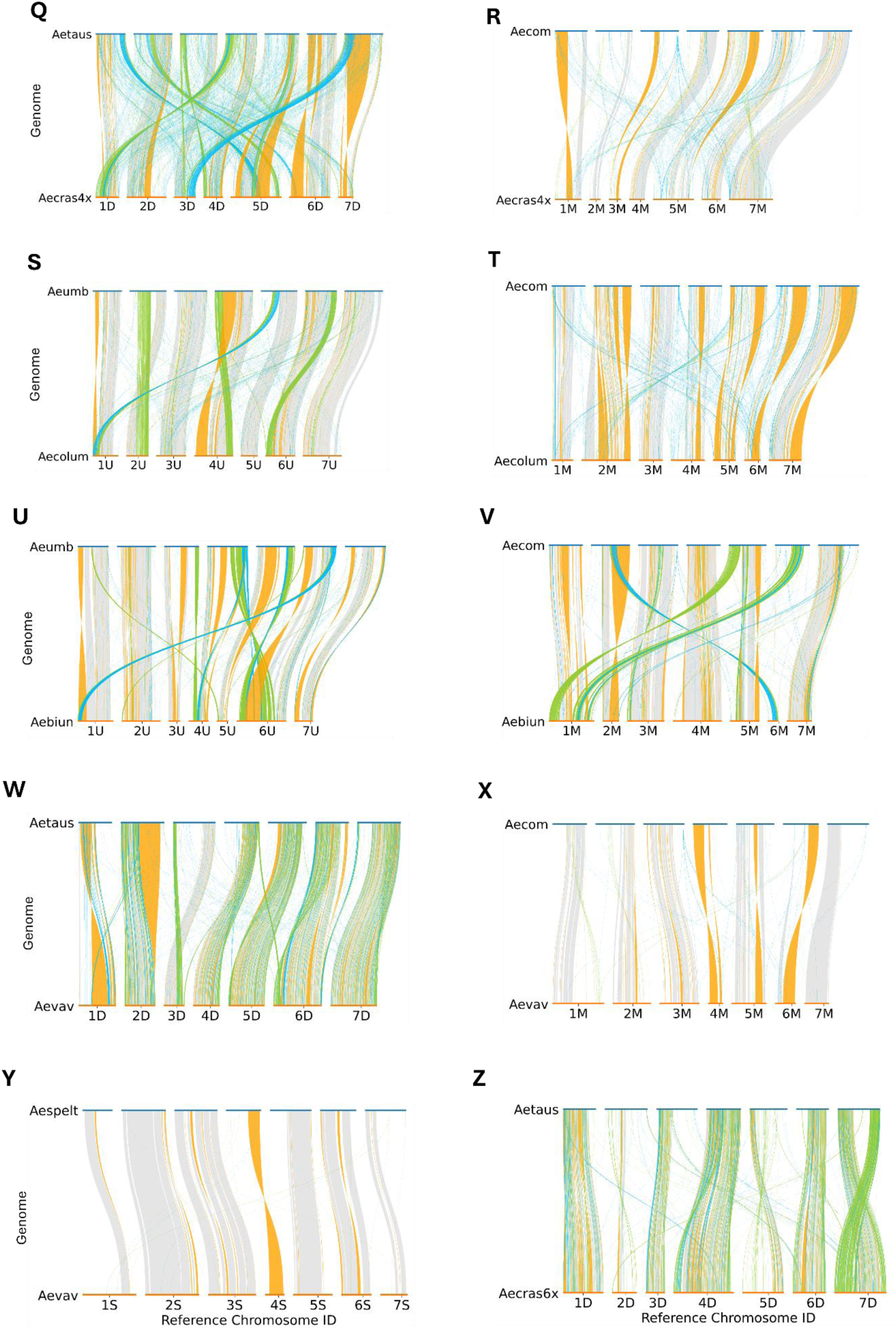
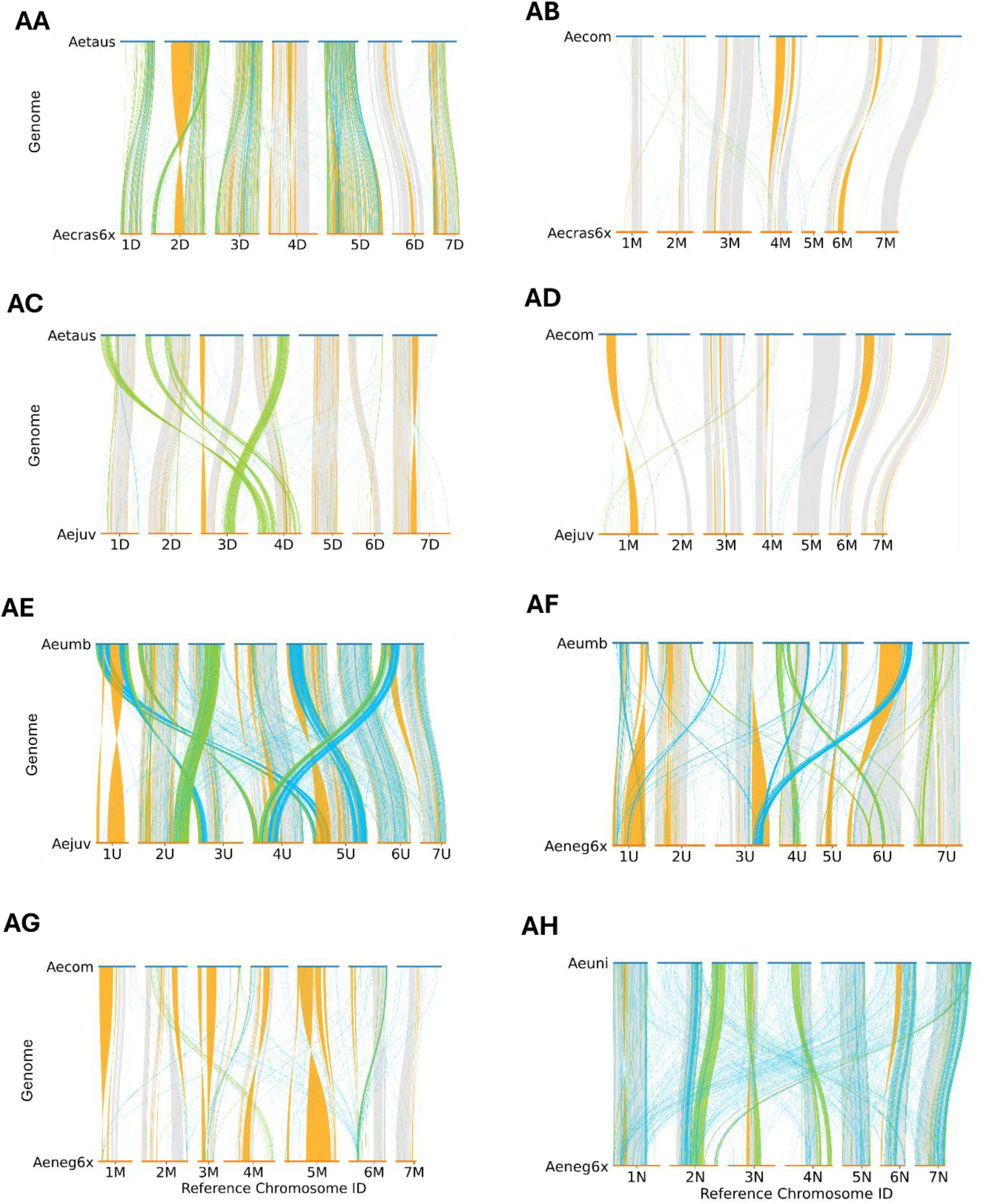
Structural variation between polyploid subgenomes and their respective diploid progenitors. **(A-AH)** Whole-genome alignments illustrating structural variation between polyploid *Aegilops* subgenomes and their corresponding diploid progenitors: **(A)** U subgenome of *Ae. neglecta* subsp. *neglecta* vs. *Ae. umbellulata*; **(B)** M subgenome of *Ae. neglecta* subsp. *neglecta* vs. *Ae. comosa*; **(C)** D subgenome of *Ae. cylindrica* vs. *Ae. tauschii*; **(D)** C subgenome of *Ae. cylindrica* vs. *Ae. markgrafii*; **(E)** D subgenome of *Ae. ventricosa* vs. *Ae. tauschii*; **(F)** N subgenome of *Ae. ventricosa* vs. *Ae. uniaristata*; **(G)** U subgenome of *Ae. triuncialis* vs. *Ae. umbellulata*; **(H)** C subgenome of *Ae. triuncialis* vs. *Ae. markgrafii*; **(I)** S subgenome of *Ae. peregrina* vs. *Ae. sharonensis*; **(J)** S subgenome of *Ae. peregrina* vs. *Ae. longissima*; **(K)** U subgenome of *Ae. peregrina* vs. *Ae. umbellulata*; **(L)** S subgenome of *Ae. kotschyi* vs. *Ae. sharonensis*; **(M)** S subgenome of *Ae. kotschyi* vs. *Ae. longissima*; **(N)** U subgenome of *Ae. kotschyi* vs. *Ae. umbellulata*; **(O)** M subgenome of *Ae. geniculata* vs. *Ae. comosa*; **(P)** U subgenome of *Ae. geniculata* vs. *Ae. umbellulata*; **(Q)** D subgenome of *Ae. crassa* subsp. *crassa* (4x) vs. *Ae. tauschii*; **(R)** M subgenome of *Ae. crassa* subsp. *crassa* (4x) vs. *Ae. comosa*; **(S)** U subgenome of *Ae. columnaris* vs. *Ae. umbellulata*; **(T)** M subgenome of *Ae. columnaris* vs. *Ae. comosa*; **(U)** U subgenome of *Ae. biuncialis* vs. *Ae. umbellulata*; **(V)** M subgenome of *Ae. biuncialis* vs. *Ae. comosa*; **(W)** D subgenome of *Ae. vavilovii* vs. *Ae. tauschii*; **(X)** M subgenome of *Ae. vavilovii* vs. *Ae. comosa*; **(Y)** S subgenome of *Ae. vavilovii* vs. *Ae. speltoides*; **(Z)** first D subgenome of *Ae. crassa* subsp. *crassa* (6x) vs. *Ae. tauschii*; **(AA)** second D subgenome of *Ae. crassa* subsp. *crassa* (6x) vs. *Ae. tauschii*; **(AB)** M subgenome of *Ae. crassa* subsp. *crassa* (6x) vs. *Ae. comosa*; **(AC)** D subgenome of *Ae. juvenalis* vs. *Ae. tauschii*; **(AD)** M subgenome of *Ae. juvenalis* vs. *Ae. comosa*; **(AE)** U subgenome of *Ae. juvenalis* vs. *Ae. umbellulata*; **(AF)** U subgenome of *Ae. neglecta* subsp. *recta* vs. *Ae. umbellulata*; **(AG)** M subgenome of *Ae. neglecta* subsp. *recta* vs. *Ae. comosa*; and **(AH)** N subgenome of *Ae. neglecta* subsp. *recta* vs. *Ae. uniaristata*. Colored links indicate syntenic regions (grey), inversions (orange), translocations (green), and duplications (blue). White regions represent unaligned or filtered sequences, which may reflect large insertions or deletions, assembly gaps, or structural variants smaller than 10 kb that are not displayed. Species abbreviations: Aeumb, *Ae. umbellulata*; Aecom, *Ae. comosa*; Aetaus, *Ae. tauschii*; Aemark, *Ae. markgrafii*; Aeuni, *Ae. uniaristata*; Aespelt, *Ae. speltoides*; Aeshar, *Ae. sharonensis*; Aelong, *Ae. longissima*; Aeneg4x, *Ae. neglecta* subsp. *neglecta*; Aecylin, *Ae. cylindrica*; Aevent, *Ae. ventricosa*; Aetriun, *Ae. triuncialis*; Aepereg, *Ae. peregrina*; Aekots, *Ae. kotschyi*; Aegen, *Ae. geniculata*; Aecras4x, *Ae. crassa* subsp. *crassa* (4x); Aecolum, *Ae. columnaris*; Aebiun, *Ae. biuncialis*; Aevav, *Ae. vavilovii*; Aecras6x, *Ae. crassa* subsp. *crassa* (6x); Aejuv, *Ae. juvenalis*; Aeneg6x, *Ae. neglecta* subsp. *recta*.

**Supplemental Figure 3.**
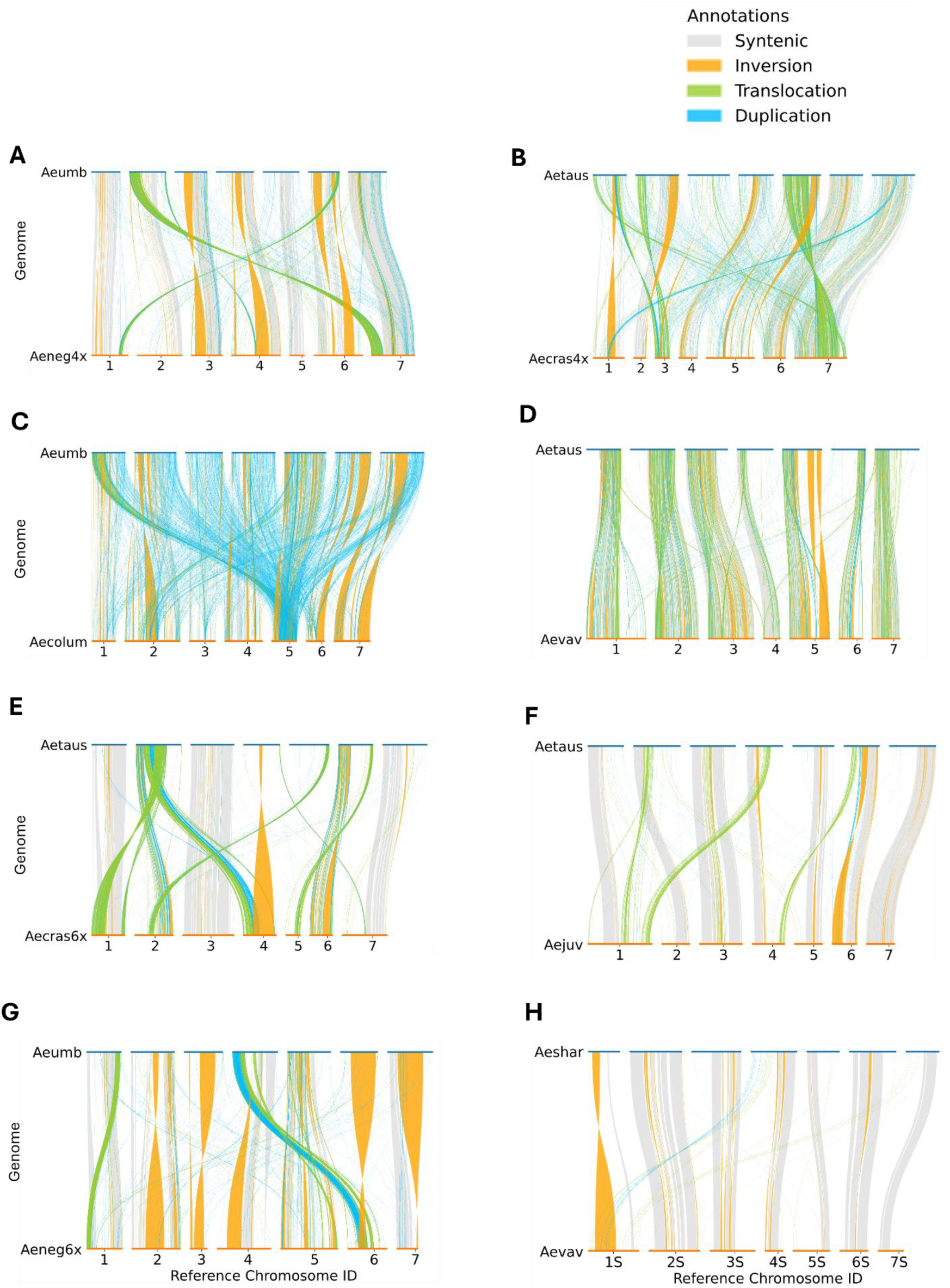
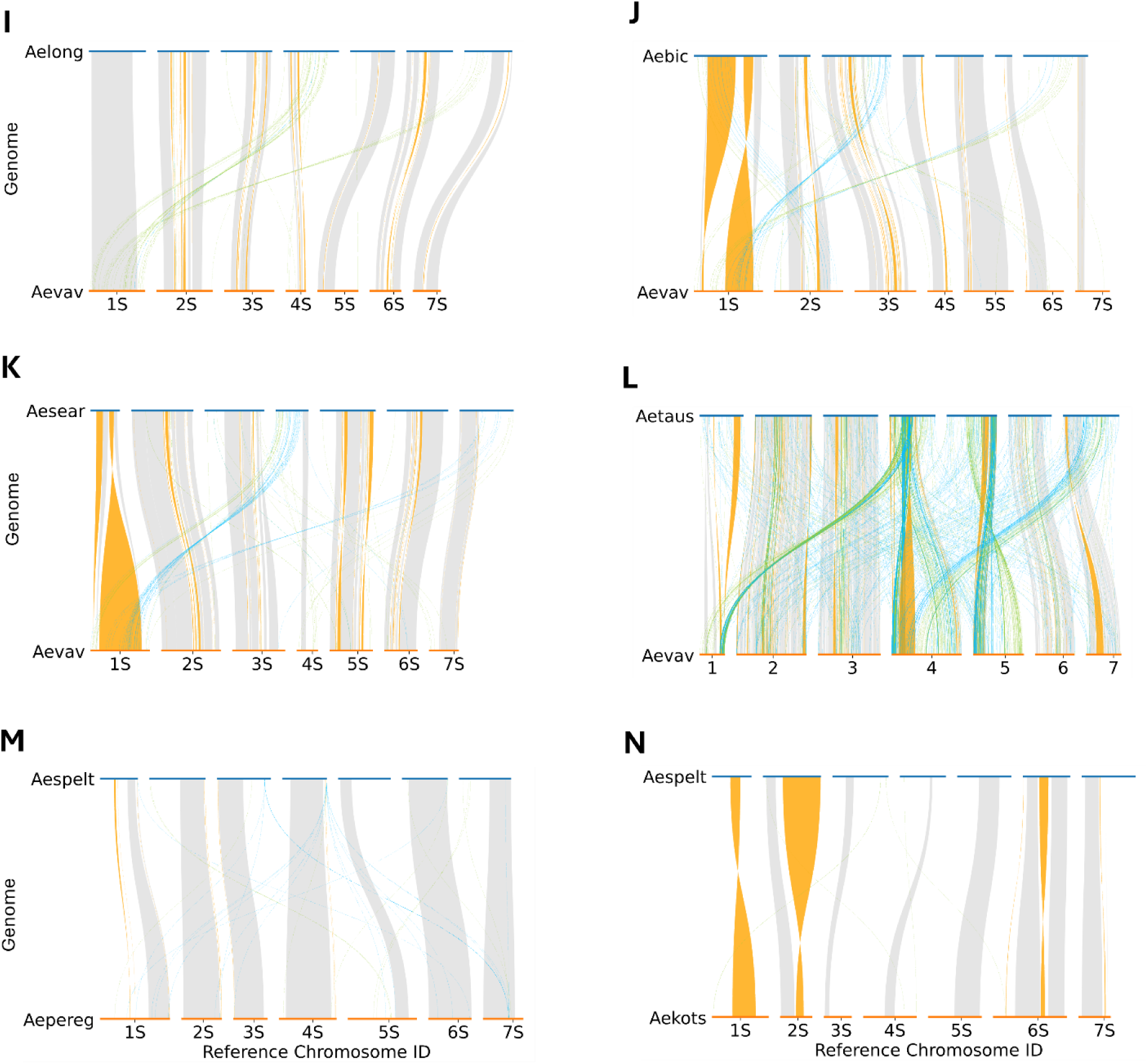
Structural variation between polyploid subgenomes and references corresponding to their coexisting subgenome counterparts. **(A–N)** Pairwise whole-genome alignments illustrating structural variation between polyploid subgenomes and references corresponding to their coexisting subgenome counterparts: **(A)** M subgenome of *Ae. neglecta* subsp. *neglecta* vs. U subgenome (*Ae. umbellulata*); **(B)** M subgenome of *Ae. crassa* subsp. *crassa* (4x) vs. D subgenome (*Ae. tauschii*); **(C)** M subgenome of *Ae. columnaris* vs. U subgenome (*Ae. umbellulata*); **(D)** M subgenome of *Ae. vavilovii* vs. D subgenome (*Ae. tauschii*); **(E)** M subgenome of *Ae. crassa* subsp. *crassa* (6x) vs. D subgenome (*Ae. tauschii*); **(F)** M subgenome of *Ae. juvenalis* vs. D subgenome (*Ae. tauschii*); **(G)** M subgenome of *Ae. neglecta* subsp. *recta* vs. U subgenome (*Ae. umbellulata*); **(H)** S subgenome of *Ae. vavilovii* vs. S^sh^ (*Ae. sharonensis*); **(I)** S subgenome of *Ae. vavilovii* vs. S^l^ (*Ae. longissima*); **(J)** S subgenome of *Ae. vavilovii* vs. S^b^ (*Ae. bicornis*); **(K)** S subgenome of *Ae. vavilovii* vs. S^s^ (*Ae. searsii*); **(L)** S subgenome of *Ae. vavilovii* vs. D subgenome (*Ae. tauschii*); **(M)** S subgenome of *Ae. peregrina* vs. S (*Ae. speltoides*); and **(N)** S subgenome of *Ae. kotschyi* vs. S (*Ae. speltoides*). Colored links indicate syntenic regions (grey), inversions (orange), translocations (green), and duplications (blue). White regions represent unaligned or filtered sequences, which may reflect large insertions or deletions, assembly gaps, or structural variants smaller than 10 kb that are not displayed. Species abbreviations: Aeumb, *Ae. umbellulata*; Aeneg4x, *Ae. neglecta* subsp. *neglecta*; Aetaus, *Ae. tauschii*; Aecras4x, *Ae. crassa* subsp. *crassa* (4x); Aecolum, *Ae. columnaris*; Aevav, *Ae. vavilovii*; *Ae. crassa* subsp. *crassa* (6x); Aejuv, *Ae. juvenalis*; Aeneg6x, *Ae. neglecta* subsp. *recta*; Aeshar, *Ae. sharonensis*; Aelong, *Ae. longissima*; Aebic, *Ae. bicornis*; Aesear, *Ae. searsii*; Aespelt, *Ae. speltoides*; Aepereg, *Ae. peregrina*; Aekots, *Ae. kotschyi*.

**Supplemental Figure 4.**
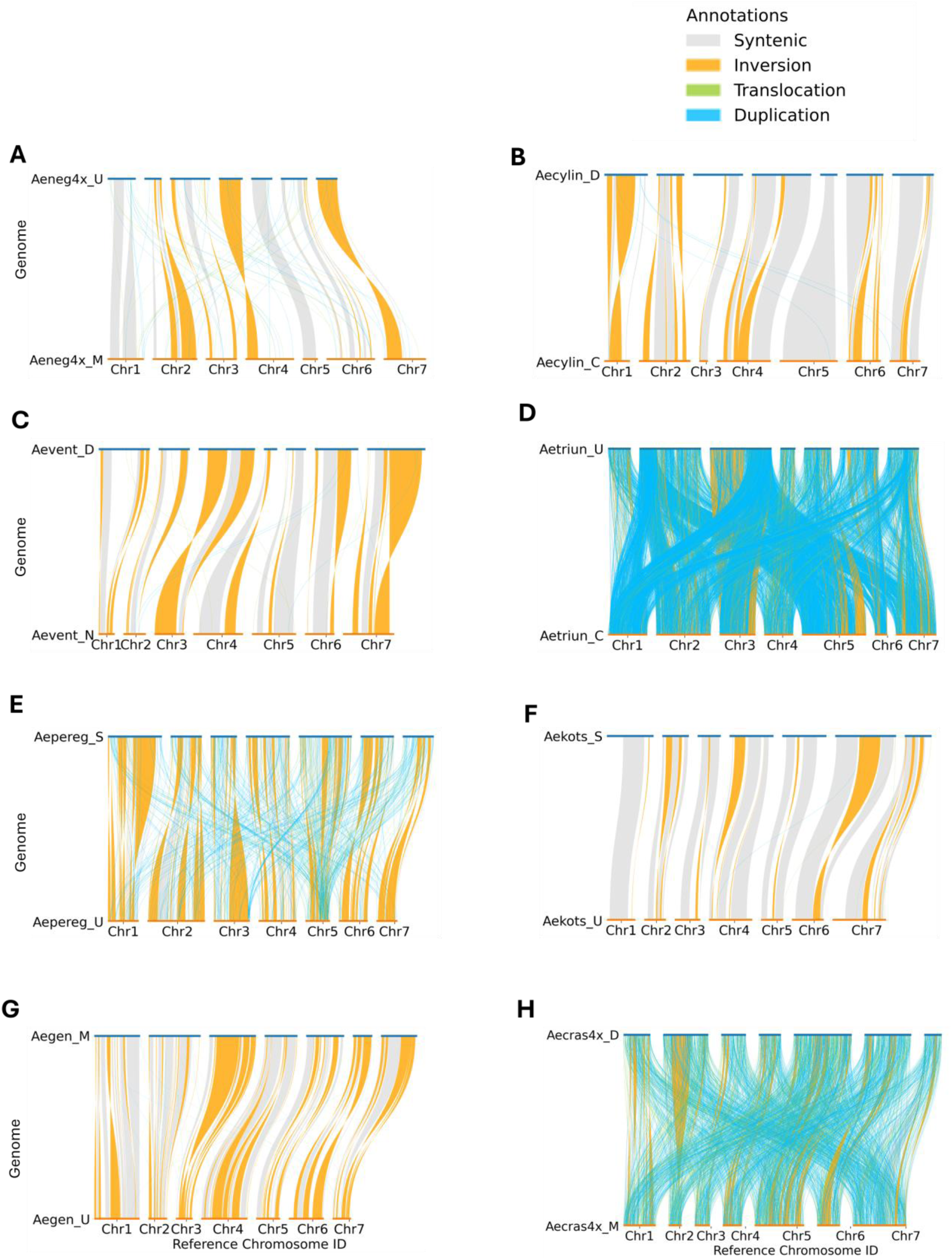
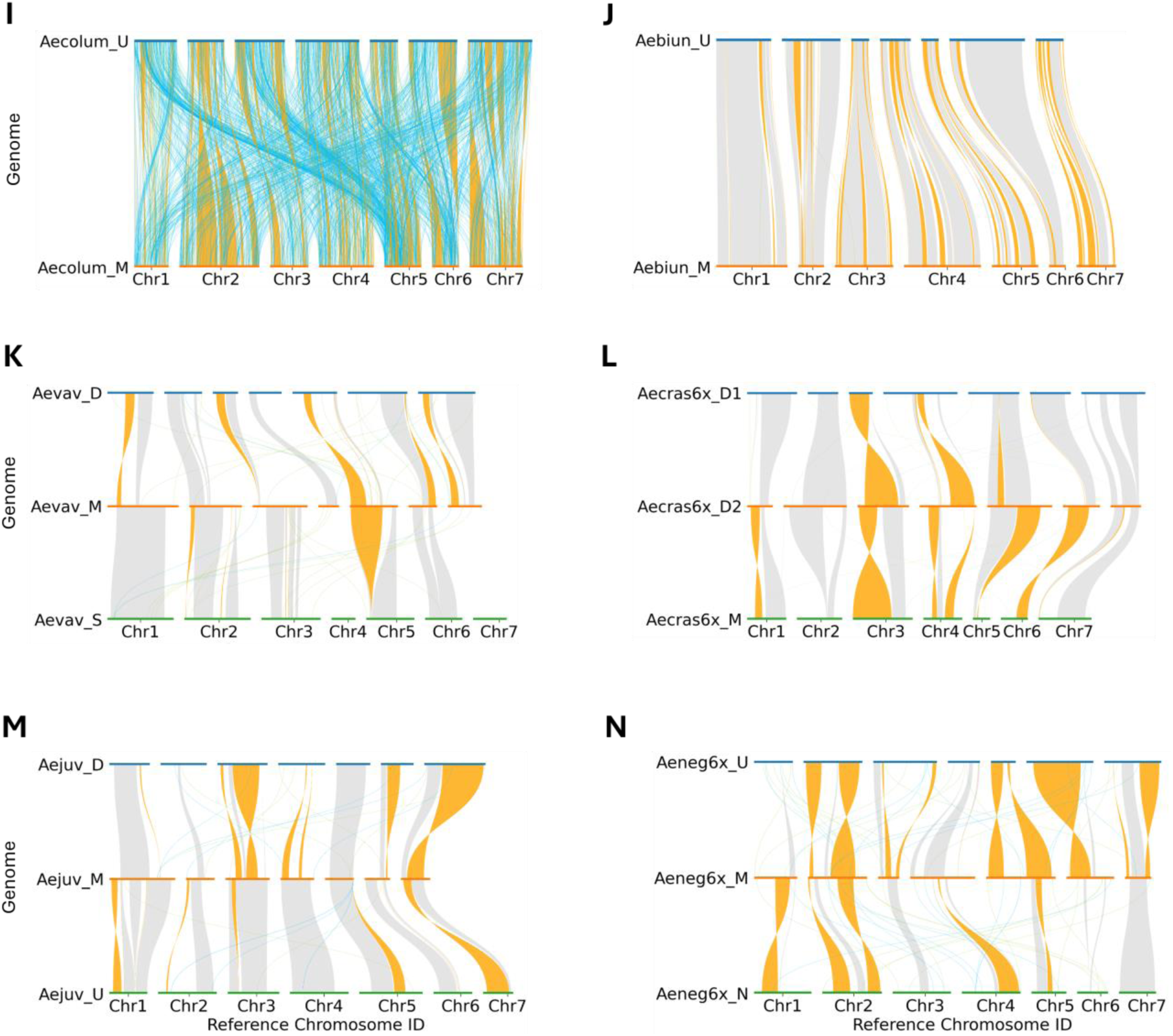
Structural variation between subgenomes within polyploid *Aegilops* species. **(A–N)** Pairwise whole-genome alignments illustrating structural variation between subgenomes within polyploid *Aegilops* species: **(A)** U vs M subgenome of *Ae. neglecta* subsp. *neglecta*; **(B)** D vs C subgenome of *Ae. cylindrica*; **(C)** D vs N subgenome of *Ae. ventricosa*; **(D)** U vs C subgenome of *Ae. triuncialis*; **(E)** S vs U subgenome of *Ae. peregrina*; **(F)** S vs U subgenome of *Ae. kotschyi*; **(G)** M vs U subgenome of *Ae. geniculata*; **(H)** D vs M subgenome of *Ae. crassa* subsp. *crassa* (4x); **(I)** U vs M subgenome of *Ae. columnaris*; **(J)** U vs M subgenome of *Ae. biuncialis*; **(K)** D vs M vs S subgenome of *Ae. vavilovii*; **(L)** D vs D vs M subgenome of *Ae. crassa* subsp. *crassa* (6x); **(M)** D vs M vs U subgenomes of *Ae. juvenalis*; **(N)** U vs M vs N subgenome of *Ae. neglecta* subsp. *recta*. Colored links indicate syntenic regions (grey), inversions (orange), translocations (green), and duplications (blue). White regions represent unaligned or filtered sequences, which may reflect large insertions or deletions, assembly gaps, or structural variants smaller than 10 kb that are not displayed. Species abbreviations: Aeneg4x, *Ae. neglecta* subsp. *neglecta*; Aecylin, *Ae. cylindrica*; Aevent, *Ae. ventricosa*; Aetriun, *Ae. triuncialis*; Aepereg, *Ae. peregrina*; Aekots, *Ae. kotschyi*; Aegen, *Ae. geniculata*; Aecras4x, *Ae. crassa* subsp. *crassa* (4x); Aecolum, *Ae. columnaris*; Aebiun, *Ae. biuncialis*; Aevav, *Ae. vavilovii*; Aecras6x, *Ae. crassa* subsp. *crassa* (6x); Aejuv, *Ae. juvenalis*; Aeneg6x, *Ae. neglecta* subsp. *recta*.

**Supplemental Figure 5.**
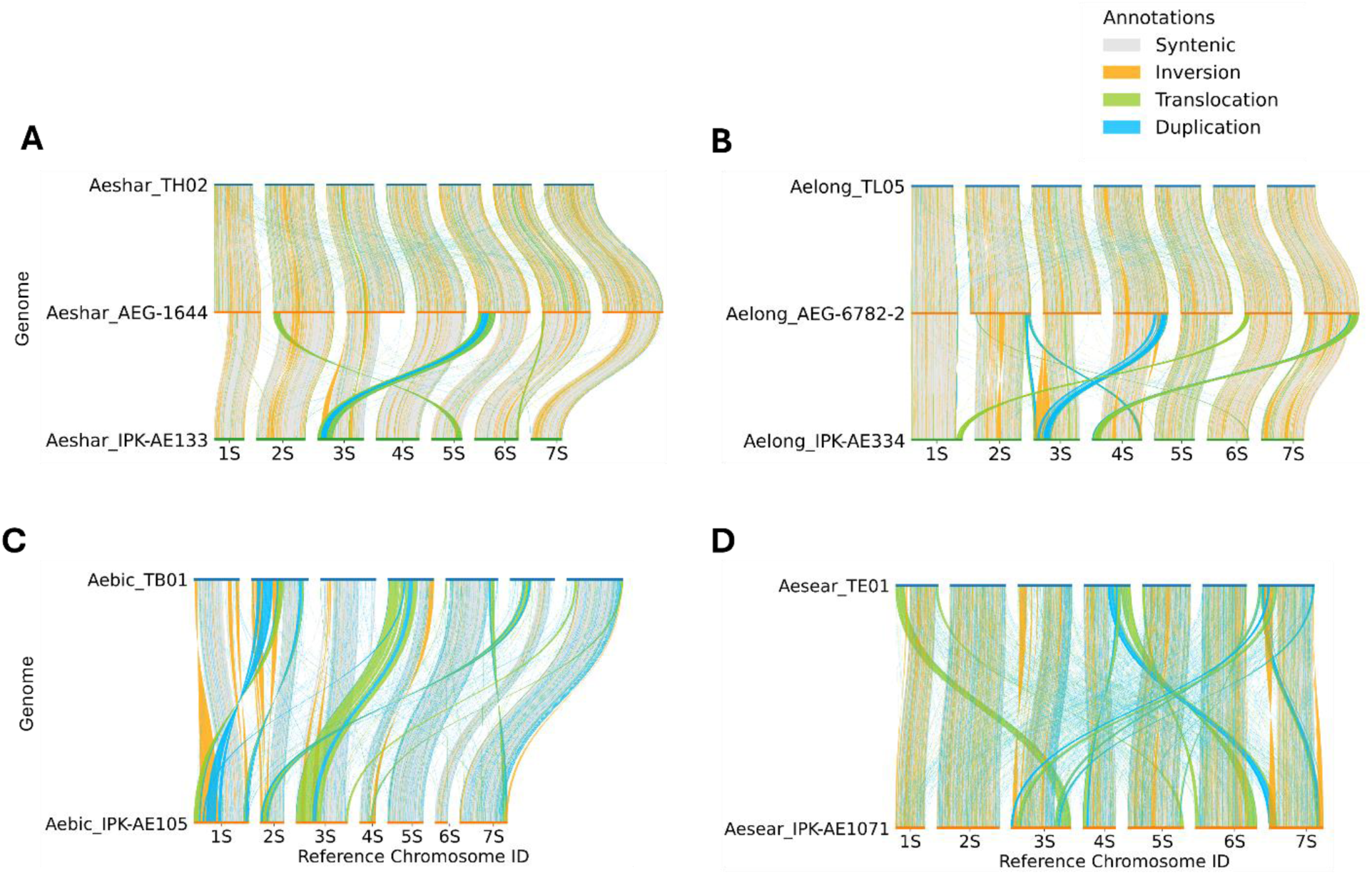
Structural variation identified in intraspecific diploid *Aegilops* genome comparisons. **(A–D)** Pairwise genome alignments illustrating structural variation between assemblies of *Ae. sharonensis*, *Ae. longissima*, *Ae. bicornis*, and *Ae. searsii*. Assemblies include previously published genomes from Li et al. (2022): Aeshar_TH02 (*Ae. sharonensis*), Aelong_TL05 (*Ae. longissima*), Aebic_TB01 (*Ae. bicornis*), and Aesear_TE01 (*Ae. searsii*); and Avni et al. (2022): Aeshar_AEG-1644 (*Ae. sharonensis*) and Aelong_AEG-6782-2 (*Ae. longissima*). Assemblies generated in this study include Aeshar_IPK-AE133 (*Ae. sharonensis*), Aelong_IPK-AE334 (*Ae. longissima*), Aebic_IPK-AE105 (*Ae. bicornis*), and Aesear_IPK-AE1071 (*Ae. searsii*). Colored links indicate syntenic regions (grey), inversions (orange), translocations (green), and duplications (blue). White regions represent unaligned or filtered sequences, which may correspond to large insertions or deletions, assembly gaps, or structural variants smaller than 10 kb that are not displayed.

## References

Abrouk, M., Wang, Y., Cavalet-Giorsa, E., Troukhan, M., Kravchuk, M., and Krattinger, S.G. (2023). Chromosome-scale assembly of the wild wheat relative *Aegilops umbellulata*. Sci. Data 10:739.

Ahmed, H.I., Heuberger, M., Schoen, A., Koo, D.-H., Quiroz-Chavez, J., Adhikari, L., Raupp, J., Cauet, S., Rodde, N., Cravero, C., et al. (2023). Einkorn genomics sheds light on history of the oldest domesticated wheat. Nature 620:830–838.

Avni, R., Lux, T., Minz-Dub, A., Millet, E., Sela, H., Distelfeld, A., Deek, J., Yu, G., Steuernagel, B., Pozniak, C., et al. (2022). Genome sequences of three *Aegilops* species of the section Sitopsis reveal phylogenetic relationships and provide resources for wheat improvement. Plant J. 110:179–192.

Avni, R., Nave, M., Barad, O., Baruch, K., Twardziok, S.O., Gundlach, H., Hale, I., Mascher, M., Spannagl, M., Wiebe, K., et al. (2017). Wild emmer genome architecture and diversity elucidate wheat evolution and domestication. Science (New York, N.Y.) 357:93–97.

Badaeva, E., Amosova, A., Samatadze, T., Zoshchuk, S., Shostak, N., Chikida, N., Zelenin, A., Raupp, W., Friebe, B., and Gill, B. (2004). Genome differentiation in *Aegilops*. 4. Evolution of the U-genome cluster. Plant System. Evol. 246:45–76.

Badaeva, E.D., Chikida, N.N., Belousova, M.K., Ruban, A.S., Surzhikov, S.A., and Zoshchuk, S.A. (2021). A new insight on the evolution of polyploid *Aegilops* species from the complex Crassa: Molecular-cytogenetic analysis. Plant System. Evol. 307:3.

Beier, S., Himmelbach, A., Colmsee, C., Zhang, X.-Q., Barrero, R.A., Zhang, Q., Li, L., Bayer, M., Bolser, D., Taudien, S., et al. (2017). Construction of a map-based reference genome sequence for barley, *Hordeum vulgare* L. Sci. Data 4:170044.

Bennetzen, J.L., and Wang, H. (2014). The contributions of transposable elements to the structure, function, and evolution of plant genomes. Ann. Rev. Plant Biol. 65:505–530.

Bernhardt, N., Brassac, J., Dong, X., Willing, E.M., Poskar, C.H., Kilian, B., and Blattner, F.R. (2020). Genome-wide sequence information reveals recurrent hybridization among diploid wheat wild relatives. Plant J. 102:493–506.

Boeckmann, B., Bairoch, A., Apweiler, R., Blatter, M.C., Estreicher, A., Gasteiger, E., Martin, M.J., Michoud, K., O’Donovan, C., Phan, I., et al. (2003). The SWISS-PROT protein knowledgebase and its supplement TrEMBL in 2003. Nucleic Acids Res. 31:365–370.

Castresana, J. (2000). Selection of conserved blocks from multiple alignments for their use in phylogenetic analysis. Mol. Biol. Evol. 17:540–552.

Cavanagh, C.R., Chao, S., Wang, S., Huang, B.E., Stephen, S., Kiani, S., Forrest, K., Saintenac, C., Brown-Guedira, G.L., Akhunova, A., et al. (2013). Genome-wide comparative diversity uncovers multiple targets of selection for improvement in hexaploid wheat landraces and cultivars. Proc. Natl. Acad. Sci. USA. 110:8057–8062.

Charmet, G. (2011). Wheat domestication: lessons for the future. C. R. Biol. 334:212–220.

Chen, N., Chen, W.-J., Yan, H., Wang, Y., Kang, H.-Y., Zhang, H.-Q., Zhou, Y.-H., Sun, G.-L., Sha, L.-N., and Fan, X. (2020). Evolutionary patterns of plastome uncover diploid-polyploid maternal relationships in Triticeae. Mol. Phylogenet. Evol. 149:106838.

Chen, Y., Zhang, Y., Wang, A.Y., Gao, M., and Chong, Z. (2021). Accurate long-read *de novo* assembly evaluation with Inspector. Genome Biol. 22:312.

Cheng, F., Wu, J., Cai, X., Liang, J., Freeling, M., and Wang, X. (2018). Gene retention, fractionation and subgenome differences in polyploid plants. Nat. Plants 4:258–268.

Cheng, H., Concepcion, G.T., Feng, X., Zhang, H., and Li, H. (2021). Haplotype-resolved de novo assembly using phased assembly graphs with hifiasm. Nature Methods 18:170–175.

Colmer, T.D., Flowers, T.J., and Munns, R. (2006). Use of wild relatives to improve salt tolerance in wheat. J. Exp. Bot. 57:1059–1078.

de Tomás, C., and Vicient, C.M. (2024). The genomic shock hypothesis: genetic and epigenetic alterations of transposable elements after interspecific hybridization in plants. Epigenomes 8:2.

Dempewolf, H., Baute, G., Anderson, J., Kilian, B., Smith, C., and Guarino, L. (2017). Past and future use of wild relatives in crop breeding. Crop Sci. 57:1070-1082.

Dos Reis, M., and Yang, Z. (2019). Bayesian molecular clock dating using genome-scale datasets. Methods Mol. Biol. 1910:309–330.

Dubcovsky, J., and Dvorak, J. (1995). Genome identification of the *Triticum crassum* complex (Poaceae) with the restriction patterns of repeated nucleotide sequences. Am. J. Bot. 82:131–140.

Dubcovsky, J., and Dvorak, J. (2007). Genome plasticity a key factor in the success of polyploid wheat under domestication. Science (New York, N.Y.) 316:1862–1866.

Dvorak, J., Terlizzi, P., Zhang, H.B., and Resta, P. (1993). The evolution of polyploid wheats: identification of the A genome donor species. Genome 36:21–31.

Dvorak, J., Luo, M.C., Yang, Z.L., and Zhang, H.B. (1998). The structure of the *Aegilops tauschii* genepool and the evolution of hexaploid wheat. Theor. Appl. Genet. 97:657–670.

Eilam, T., Anikster, Y., Millet, E., Manisterski, J., and Feldman, M. (2008). Nuclear DNA amount and genome downsizing in natural and synthetic allopolyploids of the genera *Aegilops* and *Triticum*. Genome 51:616–627.

Eilam, T., Anikster, Y., Millet, E., Manisterski, J., Sagi-Assif, O., and Feldman, M. (2007). Genome size and genome evolution in diploid Triticeae species. Genome 50:1029–1037.

Emms, D.M., and Kelly, S. (2019). OrthoFinder: phylogenetic orthology inference for comparative genomics. Genome Biol. 20:238.

Fedoroff, N. (2000). Transposons and genome evolution in plants. Proc. Natl. Acad. Sci. USA 97:7002–7007.

Feldman, M., and Levy, A.A. (2012). Genome evolution due to allopolyploidization in wheat. Genetics 192:763–774.

Feschotte, C., Jiang, N., and Wessler, S.R. (2002). Plant transposable elements: where genetics meets genomics. Nat. Rev. Genet. 3:329–341.

Flagel, L.E., and Wendel, J.F. (2009). Gene duplication and evolutionary novelty in plants. New Phytol. 183:557–564.

Friebe, B. (1996). Chromosome banding and genome analysis in diploid and cultivated polyploid wheats. In Methods of genome analysis in plants, P.P. Jauhar, ed. (CRC Press: Boca Raton, FL), pp. 39–60.

Gaut, B.S. (2002). Evolutionary dynamics of grass genomes. New Phytologist 154:15–28.

Gill, B., and Friebe, B. (2002). Cytogenetics, phylogeny and evolution of cultivated wheats. Bread wheat: improvement and production. Food and Agriculture Organization of the United Nations, Rome:71–88.

Glémin, S., Scornavacca, C., Dainat, J., Burgarella, C., Viader, V., Ardisson, M., Sarah, G., Santoni, S., David, J., and Ranwez, V. (2019). Pervasive hybridizations in the history of wheat relatives. Science Adv. 5:eaav9188.

Goel, M., and Schneeberger, K. (2022). plotsr: visualizing structural similarities and rearrangements between multiple genomes. Bioinformatics (Oxford, England) 38:2922–2926.

Goel, M., Sun, H., Jiao, W.-B., and Schneeberger, K. (2019). SyRI: finding genomic rearrangements and local sequence differences from whole-genome assemblies. Genome Biol. 20:277.

Grewal, S., Yang, C.-y., Scholefield, D., Ashling, S., Ghosh, S., Swarbreck, D., Collins, J., Yao, E., Sen, T.Z., Wilson, M., et al. (2024). Chromosome-scale genome assembly of bread wheat’s wild relative *Triticum timopheevii*. Sci. Data 11:420.

Grewal, S., Yang, C.-y., Krasheninnikova, K., Collins, J., Wood, J.M.D., Ashling, S., Scholefield, D., Kaithakottil, G.G., Swarbreck, D., Yao, E., et al. (2025). Chromosome-level haplotype-resolved genome assembly of bread wheat’s wild relative *Aegilops mutica*. Sci. Data 12:438.

Gustafsson, Ö., Kruså, M., Zencak, Z., Sheesley, R.J., Granat, L., Engström, E., Praveen, P.S., Rao, P.S.P., Leck, C., and Rodhe, H. (2009). Brown Clouds over South Asia: Biomass or Fossil Fuel Combustion? Science (New York, N.Y.) 323:495–498.

Helguera, M., Khan, I.A., Kolmer, J., Lijavetzky, D., Zhong-qi, L., and Dubcovsky, J. (2003). PCR assays for the *Lr37-Yr17-Sr38* cluster of rust resistance genes and their use to develop isogenic hard red spring wheat lines. Crop Sci. 43:1839–1847.

IWGSC (2021). Optical maps refine the bread wheat *Triticum aestivum* cv. Chinese Spring genome assembly. Plant J. 107:303–314.

Jewell, D. (1979). Chromosome banding in *Triticum aestivum* cv. Chinese Spring and *Aegilops variabilis*. Chromosoma 71:129–134.

Jewell, D., and Driscoll, C. (1983). The addition of *Aegilops variabilis* chromosomes to *Triticum aestivum* and their identification. Can. J. Genet. Cytol. 25:76–84.

Katoh, K., and Standley, D.M. (2013). MAFFT multiple sequence alignment software version 7: improvements in performance and usability. Mol. Biol. Evol. 30:772–780.

Kihara, H. (1944). Discovery of the DD-Analyser, one of the ancestors of *Triticum vulgare*. Agric. Hortic. 19:13–14.

Kilian, B., Mammen, K., Millet, E., Sharma, R., Graner, A., Salamini, F., Hammer, K., and Ozkan, H. (2011). *Aegilops* L. In Wild crop relatives: genomic and breeding resources, cereals, C. Kole, ed. (Springer: Berlin, Heidelberg), pp. 1–76.

Kozlov, A.M., Darriba, D., Flouri, T., Morel, B., and Stamatakis, A. (2019). RAxML-NG: a fast, scalable and user-friendly tool for maximum likelihood phylogenetic inference. Bioinformatics (Oxford, England) 35:4453–4455.

Kroupin, P.Y., Badaeva, E.D., Sokolova, V.M., Chikida, N.N., Belousova, M.K., Surzhikov, S.A., Nikitina, E.A., Kocheshkova, A.A., Ulyanov, D.S., and Ermolaev, A.S. (2022). *Aegilops crassa* Boiss. repeatome characterized using low-coverage NGS as a source of new FISH markers: Application in phylogenetic studies of the *Triticeae*. Front. Plant Sci. 13:980764.

Kundu, R., Casey, J., and Sung, W.-K. (2019). HyPo: super fast & accurate polisher for long read genome assemblies. bioRxiv:2019.2012.2019.882506.

Leitch, I.J., and Bennett, M.D. (2004). Genome downsizing in polyploid plants. Biol. J. Linn. Soc. 82:651–663.

Li, H., Rehman, S.u., Song, R., Qiao, L., Hao, X., Zhang, J., Li, K., Hou, L., Hu, W., Wang, L., et al. (2024). Chromosome-scale assembly and annotation of the wild wheat relative *Aegilops comosa*. Sci. Data 11:1454.

Li, L.F., Zhang, Z.B., Wang, Z.H., Li, N., Sha, Y., Wang, X.F., Ding, N., Li, Y., Zhao, J., Wu, Y., et al. (2022). Genome sequences of five Sitopsis species of *Aegilops* and the origin of polyploid wheat B subgenome. Mol. Plant 15:488–503.

Lilienfeld, F., and Kihara, H. (1934). Genomanalyse bei *Triticum* und *Aegilops*. Cytologia 6:87–122.

Lilienfeld, F.A. (1951). H. Kihara: Genome-analysis in *Triticum* and *Aegilops*. X Concluding review. Cytologia 16:101–123.

Ling, H.-Q., Ma, B., Shi, X., Liu, H., Dong, L., Sun, H., Cao, Y., Gao, Q., Zheng, S., Li, Y., et al. (2018). Genome sequence of the progenitor of wheat A subgenome *Triticum urartu*. Nature 557:424–428.

Lisch, D. (2013). How important are transposons for plant evolution? Nat. Rev. Genet. 14:49–61.

Liu, H., Wu, S., Li, A., and Ruan, J. (2021). SMARTdenovo: a *de novo* assembler using long noisy reads. GigaByte 2021:gigabyte15.

Liu, Z., Yang, F., Wan, H., Deng, C., Hu, W., Fan, X., Wang, J., Yang, M., Feng, J., Wang, Q., et al. (2025). Genome architecture of the allotetraploid wild grass *Aegilops ventricosa* reveals its evolutionary history and contributions to wheat improvement. Plant Commun. 6:101131.

Luo, M.-C., Gu, Y.Q., Puiu, D., Wang, H., Twardziok, S.O., Deal, K.R., Huo, N., Zhu, T., Wang, L., Wang, Y., et al. (2017). Genome sequence of the progenitor of the wheat D genome *Aegilops tauschii*. Nature 551:498–502.

Maccaferri, M., Harris, N.S., Twardziok, S.O., Pasam, R.K., Gundlach, H., Spannagl, M., Ormanbekova, D., Lux, T., Prade, V.M., Milner, S.G., et al. (2019). Durum wheat genome highlights past domestication signatures and future improvement targets. Nat. Genet. 51:885–895.

Manni, M., Berkeley, M.R., Seppey, M., Simão, F.A., and Zdobnov, E.M. (2021). BUSCO update: novel and streamlined workflows along with broader and deeper phylogenetic coverage for scoring of eukaryotic, prokaryotic, and viral genomes. Mol. Biol. Evol. 38:4647–4654.

Marçais, G., Delcher, A.L., Phillippy, A.M., Coston, R., Salzberg, S.L., and Zimin, A. (2018). MUMmer4: A fast and versatile genome alignment system. PLoS Comput. Biol. 14:e1005944.

Marcussen, T., Sandve, S.R., Heier, L., Spannagl, M., Pfeifer, M., Jakobsen, K.S., Wulff, B.B., Steuernagel, B., Mayer, K.F., and Olsen, O.A. (2014). Ancient hybridizations among the ancestral genomes of bread wheat. Science (New York, N.Y.) 345:1250092.

McFadden, E., and Sears, E. (1946). The origin of *Triticum spelta* and its free-threshing hexaploid relatives. J. Hered. 37:81–107.

Neumann, P., Novák, P., Hoštáková, N., and Macas, J. (2019). Systematic survey of plant LTR-retrotransposons elucidates phylogenetic relationships of their polyprotein domains and provides a reference for element classification. Mob. DNA 10:1.

Nevo, E., and Chen, G. (2010). Drought and salt tolerances in wild relatives for wheat and barley improvement. Plant Cell Environ. 33:670–685.

Olivera, P.D., Rouse, M.N., and Jin, Y. (2018). Identification of new sources of resistance to wheat stem rust in *Aegilops* spp. in the tertiary genepool of wheat. Front. Plant Sci. 9:1719.

Ou, S., Su, W., Liao, Y., Chougule, K., Agda, J.R.A., Hellinga, A.J., Lugo, C.S.B., Elliott, T.A., Ware, D., Peterson, T., et al. (2019). Benchmarking transposable element annotation methods for creation of a streamlined, comprehensive pipeline. Genome Biol. 20:275.

Padmarasu, S., Himmelbach, A., Mascher, M., and Stein, N. (2019). In situ Hi-C for plants: an improved method to detect long-range chromatin interactions. Methods Mol. Biol. 1933:441–472.

Parisod, C., Alix, K., Just, J., Petit, M., Sarilar, V., Mhiri, C., Ainouche, M., Chalhoub, B., and Grandbastien, M.A. (2010). Impact of transposable elements on the organization and function of allopolyploid genomes. New Phytol. 186:37–45.

Petersen, G., Seberg, O., Yde, M., and Berthelsen, K. (2006). Phylogenetic relationships of *Triticum* and *Aegilops* and evidence for the origin of the A, B, and D genomes of common wheat (*Triticum aestivum*). Mol. Phylogenet. Evol. 39:70–82.

Pulido, M., and Casacuberta, J.M. (2023). Transposable element evolution in plant genome ecosystems. Curr. Opin. Plant Biol. 75:102418.

Salamini, F., Özkan, H., Brandolini, A., Schäfer-Pregl, R., and Martin, W. (2002). Genetics and geography of wild cereal domestication in the near east. Nat. Rev. Genet. 3:429–441.

Schneider, A., Molnár, I., and Molnár-Láng, M. (2008). Utilisation of *Aegilops* (goatgrass) species to widen the genetic diversity of cultivated wheat. Euphytica 163:1–19.

Stiehler, F., Steinborn, M., Scholz, S., Dey, D., Weber, A.P.M., and Denton, A.K. (2020). Helixer: cross-species gene annotation of large eukaryotic genomes using deep learning. Bioinformatics (Oxford, England) 36:5291–5298.

Suyama, M., Torrents, D., and Bork, P. (2006). PAL2NAL: robust conversion of protein sequence alignments into the corresponding codon alignments. Nucleic Acids Res. 34:W609–612.

Tadesse, W., Sanchez-Garcia, M., Assefa, S.G., Amri, A., Bishaw, Z., Ogbonnaya, F., and Baum, M. (2019). Genetic gains in wheat breeding and its role in feeding the world. Crop Breed. Genet. Genom. 1:e190005.

Van Slageren, M. (1994). Wild wheats: a monograph of Aegilops L. and *Amblyopyrum* (Jaub. & Spach) Eig (*Poaceae*) (Agricultural University Wageningen).

Vicient, C.M., and Casacuberta, J.M. (2017). Impact of transposable elements on polyploid plant genomes. Ann. Bot. 120:195–207.

Vogel, J.P., Garvin, D.F., Mockler, T.C., Schmutz, J., Rokhsar, D., Bevan, M.W., Barry, K., Lucas, S., Harmon-Smith, M., Lail, K., et al. (2010). Genome sequencing and analysis of the model grass *Brachypodium distachyon*. Nature 463:763–768.

Wang, H., Sun, S., Ge, W., Zhao, L., Hou, B., Wang, K., Lyu, Z., Chen, L., Xu, S., Guo, J., et al. (2020). Horizontal gene transfer of Fhb7 from fungus underlies Fusarium head blight resistance in wheat. Science (New York, N.Y.) 368:eaba5435.

Wang, L., Zhu, T., Rodriguez, J.C., Deal, K.R., Dubcovsky, J., McGuire, P.E., Lux, T., Spannagl, M., Mayer, K.F.X., Baldrich, P., et al. (2021). *Aegilops tauschii* genome assembly Aet v5.0 features greater sequence contiguity and improved annotation. G3 (Bethesda) 11:jkab325.

Weidner, A., Röder, M.S., and Börner, A. (2012). Mapping wheat powdery mildew resistance derived from *Aegilops markgrafii*. Plant Genet. Resour. 10:137–140.

Wicker, T., Matthews, D.E., and Keller, B. (2002). TREP: a database for *Triticeae* repetitive elements. Trends Plant Sci. 7:561–562.

Wicker, T., Sabot, F., Hua-Van, A., Bennetzen, J.L., Capy, P., Chalhoub, B., Flavell, A., Leroy, P., Morgante, M., and Panaud, O. (2007). A unified classification system for eukaryotic transposable elements. Nat. Rev. Genet. 8:973–982.

Wicker, T., Gundlach, H., Spannagl, M., Uauy, C., Borrill, P., Ramírez-González, R.H., De Oliveira, R., Mayer, K.F.X., Paux, E., Choulet, F., et al. (2018). Impact of transposable elements on genome structure and evolution in bread wheat. Genome Biol. 19:103.

Wu, T.D., and Watanabe, C.K. (2005). GMAP: a genomic mapping and alignment program for mRNA and EST sequences. Bioinformatics (Oxford, England) 21:1859–1875.

Yang, Y., Cui, L., Lu, Z., Li, G., Yang, Z., Zhao, G., Kong, C., Li, D., Chen, Y., Xie, Z., et al. (2023). Genome sequencing of *Sitopsis* species provides insights into their contribution to the B subgenome of bread wheat. Plant Commun. 4:100567.

Yu, G., Matny, O., Champouret, N., Steuernagel, B., Moscou, M.J., Hernández-Pinzón, I., Green, P., Hayta, S., Smedley, M., Harwood, W., et al. (2022). *Aegilops sharonensis* genome-assisted identification of stem rust resistance gene *Sr62*. Nat. Commun. 13:1607.

Zeng, X., Yi, Z., Zhang, X., Du, Y., Li, Y., Zhou, Z., Chen, S., Zhao, H., Yang, S., Wang, Y., et al. (2024). Chromosome-level scaffolding of haplotype-resolved assemblies using Hi-C data without reference genomes. Nat. Plants 10:1184–1200.

Zhang, H.-B., Dvořák, J., and Waines, J.G. (1992). Diploid ancestry and evolution of *Triticum kotschyi* and *T. peregrinum* examined using variation in repeated nucleotide sequences. Genome 35:182–191.

Zhang, M., Zhang, Y., Scheuring, C.F., Wu, C.-C., Dong, J.J., and Zhang, H.-B. (2012). Preparation of megabase-sized DNA from a variety of organisms using the nuclei method for advanced genomics research. Nat. Protoc. 7:467–478.

Zhang, R.-G., Li, G.-Y., Wang, X.-L., Dainat, J., Wang, Z.-X., Ou, S., and Ma, Y. (2022). TEsorter: An accurate and fast method to classify LTR-retrotransposons in plant genomes. Hort. Res. 9:uhac017.

Zhang, Z., Lv, R., Wang, B., Xun, H., Liu, B., and Xu, C. (2023). Effects of allopolyploidization and homoeologous chromosomal segment exchange on homoeolog expression in a synthetic allotetraploid wheat under variable environmental conditions. Plants 12:3111.

Zhao, L., Ning, S., Yu, J., Hao, M., Zhang, L., Yuan, Z., Zheng, Y., and Liu, D. (2016). Cytological identification of an *Aegilops variabilis* chromosome carrying stripe rust resistance in wheat. Breed. Sci. 66:522–529.

Zhou, C., McCarthy, S.A., and Durbin, R. (2022). YaHS: yet another Hi-C scaffolding tool. Bioinformatics (Oxford, England) 39:btac808.

Zhu, T., Wang, L., Rodriguez, J.C., Deal, K.R., Avni, R., Distelfeld, A., McGuire, P.E., Dvorak, J., and Luo, M.C. (2019). Improved genome sequence of wild emmer wheat Zavitan with the aid of optical maps. G3 (Bethesda) 9:619–624.

